# CMTM6 maintains B cell-intrinsic CD40 expression to regulate anti-tumor immunity

**DOI:** 10.1101/2024.03.05.583639

**Authors:** Yiru Long, Runqiu Chen, Wenlong Chang, Fanglin Li, Longhua Gu, Jianhua Sun, Jing Chen, Likun Gong

**Author notes:** Correspondence: Likun Gong.

## Abstract

**Highlights:**

- B cells participate in anti-tumor immunity by influencing the intratumoral infiltration and function of T cells;
- CMTM6 cis-interacts with CD40 and inhibits ubiquitin/proteasome-mediated CD40 degradation to maintain CD40 cell membrane levels;
- Loss of B cell-intrinsic CMTM6 significantly reduces CD40 signaling-mediated B cell activation, survival, differentiation, T/B cell interaction and anti-tumor immunity;
- B cell-intrinsic CMTM6 deficiency leads to a significant reduction in the anti-tumor activity CD40 agonists and ICB therapy.

**In Brief:** Long et al. demonstrate that B-cell intrinsic CMTM6 regulates CD40 signaling and function via maintaining the cell membrane level of B-cell CD40 through ubiquitin-proteasome pathway inhibition, thereby affecting B cell function, anti-tumor B-cell immunity and the efficacy of CD40 agonist and ICB therapy.

Graphic Abstracts

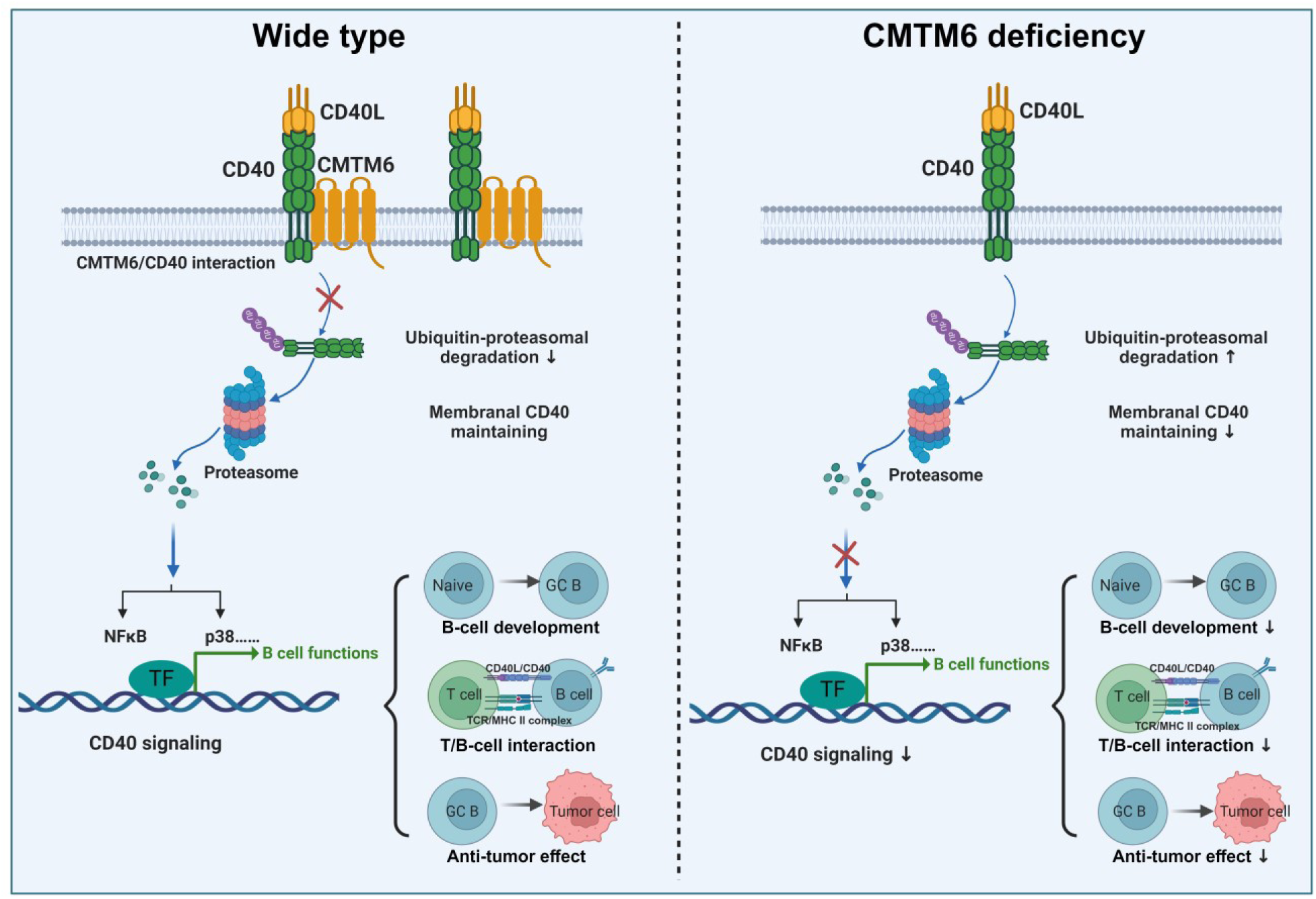

The essential role of B cells and B cell intrinsic molecules in tumor immunity is beginning to be recognized. Tumor cell CMTM6 is a novel tumor immunoregulator involved in maintaining membrane levels of several important molecules, such as PD-L1 and CD58. Host CMTM6 may also play a function in the tumor microenvironment. Here, we found that CMTM6 was highly expressed in splenic B cells and tumor-infiltrating B cells. CMTM6 deficiency resulted in impaired splenic development, germinal center B cell differentiation, memory B cell differentiation, T/B cell interaction and B cell anti-tumor immune responses. Through multi-omics data mining and B-cell agonist screening, we identified that CMTM6 interacted with CD40 and maintained CD40 membrane levels in B cells. CMTM6 cis-interacts with CD40 and inhibits ubiquitin/proteasome-mediated CD40 degradation. CMTM6 deficiency led to impaired CD40 signaling-mediated B cell activation, survival, proliferation, differentiation and T/B cell interaction. In vivo, CMTM6 deficiency leads to a significant decrease in the anti-tumor activity of ICB therapy and B cell-dependent CD40 agonists. Collectively, B-cell intrinsic CMTM6 maintains B cell CD40 levels and signaling to promote B cell function and anti-tumor immunity.

## INTRODUCTION

T cells are absolutely central to tumor immunotherapy as represented by immune checkpoint blockade (ICB) therapies, and the primary objective of these therapies is to restore or improve T cell anti-tumor activity. However, the complex composition of the tumor microenvironment (TME), as well as the current status of disappointing treatment response rates, indicate that focusing solely on T cells is inadequate^1^. T and B cells are crucial to adaptive immune responses, but the importance of B cell in antitumor immunity remains unappreciated. While B cells can eliminate tumor cells through tumor-targeting antibodies that facilitate antibody-dependent cellular cytotoxicity (ADCC) and antibody-dependent cellular phagocytosis (ADCP), previous research has mostly concentrated on the tumor-promoting impacts of B cells^2^. Studies on the critical role tumor infiltrating B-lymphocytes (TIL-B) play in the ICB response and antitumor effects have emerged in recent years. The development of intratumoral tertiary lymphoid structures (TLS) relies on TIL-B cells, which are essential for the acquisition and presentation of tumor antigens, as well as for the promotion of tumor-specific T follicular helper cells (Tfh) differentiation and anti-tumor CD8^+^ T-cell activation^3,4^. Patients with early-stage lung adenocarcinoma, melanoma, sarcoma, or renal cell carcinoma exhibit a positive correlation between B-cells/TLS and their response to cancer immunotherapy^5–8^. Besides, molecules like TIM-1 have been discovered to regulate B-cell responses to tumors, indicating potential therapeutic approaches for targeting TIL-B^9,10^.

CKLF-like MARVEL transmembrane domain-containing protein 6 (CMTM6), identified in 2003, is an M-shaped four-transmembrane protein^11^. CMTM6 has been reported to be associated with the prognosis and advancement of various clinical malignancies, and it is thought to affect tumor progression via several possible pathways, such as the WNT pathway^12–15^. In recent years, tumor cell intrinsic CMTM6 has been discovered as a novel tumor immunoregulatory protein.

CMTM6 is a new post-translational modification regulatory molecule of PD-L1, capable of inhibiting the endosomal-lysosomal and ubiquitin-proteasomal degradation pathways of PD-L1, extending the half-life of tumoral PD-L1, and modulating tumor immunity^16,17^. CMTM6 has also been shown to prevent the lysosomal degradation of CD58 in tumor cells, modulating the anti-tumor responses of T cells^18,19^. Our previous studies revealed that tumor cell CMTM6 could regulate CD8^+^ T cell anti-tumor immunity by maintaining PD-L1 bidirectional signaling and PD-L1-independent β-catenin/CCL4 signaling^20,21^. We also developed two drug candidates,

CMTM6-targeted nanobodies and adeno-associated viruses, and confirmed their significant antitumor activity in vivo^20,21^. However, the expression and function of CMTM6 in tumor-infiltrating immune cells have not been systematically studied.

In this study, we investigated the anti-tumor immune role of B cells with an eye on CMTM6, and systematically elucidated the expression of CMTM6 in B cells as well as the regulatory roles and mechanisms of CMTM6 on B cell function and anti-tumor immunity.

## RESULTS

### B cells influence the TME and anti-tumor T cell responses

We initially examined the impact of total B lymphocytes on tumor immunity. We used the anti-CD20 antibody to delete B cells in tumor-bearing mice and confirmed the effect in draining lymph nodes (dLN) and tumors at the anatomical endpoint (Figures S1A and S1B). In the B16F10 tumor model, we discovered that in the absence of B cells, the leukocytes infiltrating the tumor were dramatically reduced, particularly CD4^+^ and CD8^+^ T cells, to roughly 15% of their original levels (Figure S1C). Similarly, in the MC38 tumor model, B-cell deletion also interfered with intratumoral infiltration of leukocytes and T cells (Figure S1E). Moreover, in the draining lymph nodes, the expression of T-cell effector molecules, such as interferon-γ (IFN-γ), granzyme B (GzmB), and perforin, was impaired after B-cell deletion (Figures S1D and S1E). We also observed in an in situ intestinal tumor model, APC-min, that deletion of B cells affects tumor progression (Figure S2A). Deletion of B cells did not increase the expression of inhibitory molecules such as PD-1 and PD-L1 in T cells in mesenteric lymph nodes (MLN; Figure S2B).

Similarly, deletion of B cells decreased the expression of T cell effector molecules in the MLN (Figure S2C and S2D). These findings, once again, point to the fact that total B cells generally have antitumor effects and can encourage T cell infiltration and function within tumors.

### CMTM6 is expressed in peripheral and tumor B cells

Using open-source single-cell sequencing data^22^, we determined that CMTM6 was more abundantly expressed on B cells in the peripheral blood of humans or mice (Figure S3A). CMTM6 was similarly extremely expressed in B cells that infiltrated the tumor (Figure S3B). We also confirmed the higher levels of CMTM6 in mouse splenic B cells via flow cytometry (Figure S3C). On the basis of these results, we hypothesized that CMTM6 serves an essential role in B cells.

### CMTM6 is a potential regulator of B-cell anti-tumor immunity

We then proceeded to compare the effects of CMTM6 knockout (KO) on the antitumor activity of B cells. When comparing the differences in immune organs between wild-type (WT) and *Cmtm6* KO mice, we found that the spleen size differed significantly between the two, with CMTM6 defects resulting in mice with smaller spleens and significantly lower spleen indexes (Figure 1A). In the MC38-OVA and B16-OVA subcutaneous tumor models, the spleens of *Cmtm6* KO mice also were smaller (Figure 1B). Considering that B cells are the predominant cell type in the spleen, we hypothesized that CMTM6 may affect B cell development, proliferation, or survival.

**Figure 1.**
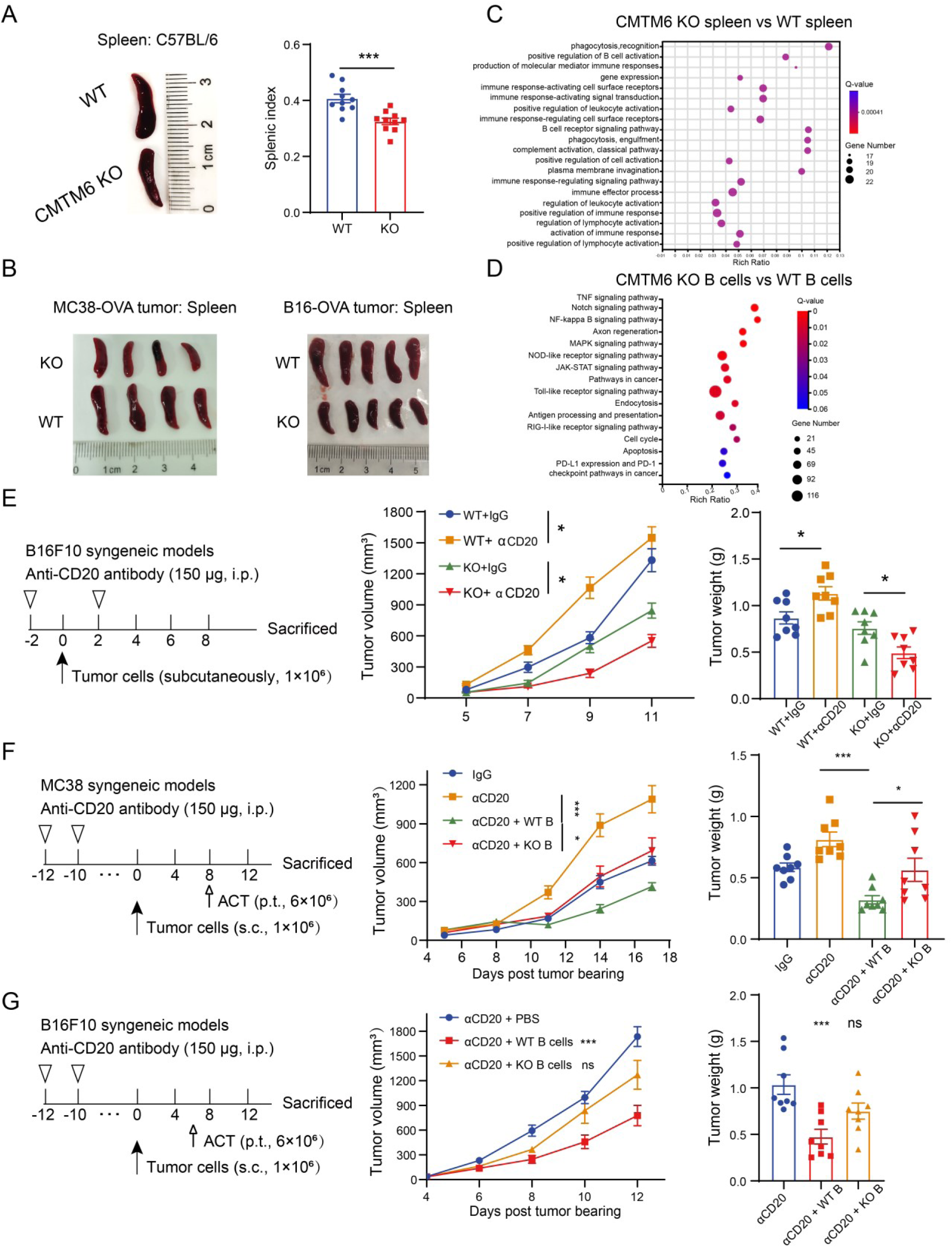
CMTM6 regulates B-cell anti-tumor immunity. (A) Representative spleen photographs and statistical analysis of splenic index (splenic weight * 100% / body weight) in WT and *Cmtm6* KO mice (n = 10). (B) Representative spleen photographs in WT and *Cmtm6* KO mice bearing MC38-OVA tumor (n = 4) or B16-OVA tumor (n = 5). (C) RNA-seq analysis of spleen tissues from WT and *Cmtm6* KO mice (n = 3). (D) RNA-seq analysis of primary splenic B cells from WT and *Cmtm6* KO mice (n = 3). (E) B16F10 tumor growth and tumor weight in WT and *Cmtm6* KO mice after B cell deletion by anti-CD20 antibody (n = 8). (F and G) Tumor growth and tumor weight of MC38 tumors (F) or B16F10 tumors (G) in mice with B cell deletion and WT or *Cmtm6* KO B cells infusion (n = 8). The data are presented as the mean ± SEM. * p < 0.05; *** p < 0.001; ns not significant by unpaired t test or one-way ANOVA followed by Tukey’s multiple comparisons test.

To investigate whether CMTM6 affects certain biological processes in B cells, we first performed RNA sequencing (RNA-seq) analysis on spleen tissues of WT and *Cmtm6* KO mice, and the results showed that the differential genes were indeed enriched in several pathways related to B cell, such as “positive regulation of B cell activation” (Figure 1C). Further, we isolated splenic B cells from WT and *Cmtm6* KO mice by magnetic cell sorting, followed by RNA-seq analysis, which showed enrichment of differential genes into immune and cancer-related pathways (Figure 1D). These results once again suggest a correlation between CMTM6 and B cell function.

In vivo, we eliminated B cells using the anti-CD20 antibody and discovered that B cell deletion resulted in faster B16F10 tumor growth in WT mice, implying that total B cells have anti-tumor effects (Figure 1E). In *Cmtm6* KO mice, the deletion of B cells further impeded tumor progression in vivo, indicating that CMTM6-deficient B cells have impaired anti-tumor activity (Figure 1E). To facilitate a more precise comparison of the antitumor effects of WT B cells and CMTM6-deficient B cells, we infused them peritumorally into mice with original B cell deletion (Figure S4A). The results indicated that, in the MC38 or B16F10 tumor model, tumor progression was inhibited by the infiltration of WT B cells with inhibition rates (IR) of 60.6% and 37.1% separately, whereas the infiltration of CMTM6-deficient B cells did not achieve noticeable antitumor effects (Figures 1F and 1G). Furthermore, the efficacy of CMTM6-deficient B cells in promoting T-cell responses was also compromised in the dLN of B16F10 tumors (Figure S4B). Taken together, these findings suggest that CMTM6 may be associated with B cell function and involved in maintaining the anti-tumor immune response of B cells.

### CMTM6 affects T/B cell interactions

Next, we planned to systematically investigate the regulatory role of CMTM6 on B cells. We first constructed B-cell CMTM6 conditional knockout mice (BKO; Figure S5). In contrast to CD19-iCre mice, B-cell CMTM6 deficiencies accelerated the progression of B16F10 and MC38 tumors (Figures 2A and 2B). Similarly, effector levels of CD4^+^ T and CD8^+^ T cells in the dLN were also lower in BKO mice (Figures 2C). Expression of inhibitory molecules in CD4^+^ T and CD8^+^ T cells in the dLN was also not affected by CMTM6 defects in B cells (Figures 2D). These data hint more directly at the importance of the anti-tumor effects of CMTM6 on B cells.

**Figure 2.**
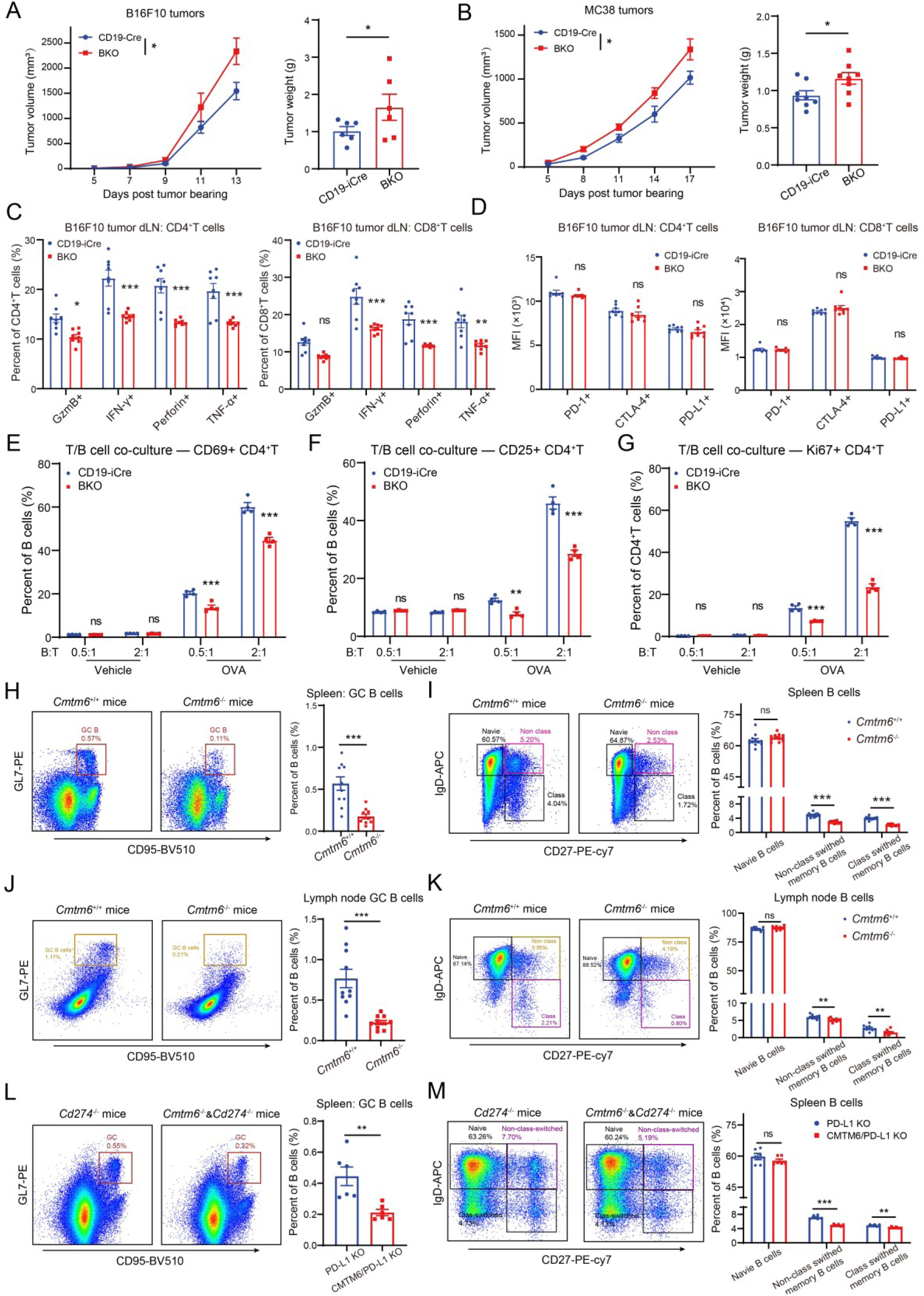
CMTM6 affects T/B cell interactions and B cell differentiation. (A and B) Tumor growth and tumor weight of B16F10 tumors (A; n = 6) or MC38 tumors (B; n = 8) in CD19-iCre or BKO mice. (C) Effector molecule levels of dLN T cells in the B16F10 tumor model (n = 8). (D) Inhibitory molecule levels of dLN T cells in the B16F10 tumor model (n = 8). (E and F) The activation levels (E: CD69^+^; F: CD25^+^) of OT-Ⅱ CD4^+^ T cell when co-cultured with CD19-iCre or BKO B cells for 24 h. (G) The proliferation levels (Ki67^+^) of OT-Ⅱ CD4^+^ T cell when co-cultured with CD19-iCre or BKO B cells for 72 h. (H and I) Representative gating strategy of splenic naïve B cells, non-class-switching memory B cells, and class-switching memory B cells, germinal center B, and statistical analysis in WT or *Cmtm6* KO mice (n = 10). (J and K) Representative gating strategy of lymph node naïve B cells, non-class-switching memory B cells, and class-switching memory B cells, germinal center B, and statistical analysis in WT or *Cmtm6* KO mice (n = 10). (L and M) Representative gating strategy of splenic naïve B cells, non-class-switching memory B cells, and class-switching memory B cells, germinal center B, and statistical analysis in *Cmtm6*/*Cd274* double-KO mice and *Cd274* KO mice (n = 10). The data are presented as the mean ± SEM. * p < 0.05; ** p < 0.01; *** p < 0.001; ns not significant by unpaired t test or two-way ANOVA followed by Tukey’s multiple comparisons test.

Given the implications for T-cell effects in the aforementioned data, our initial emphasis was on the impact of CMTM6 on T/B cell interactions. In a co-culture model of OT-II CD4^+^ T cells and B cells, OVA-loaded B cells effectively promoted CD4^+^ T activation and proliferation (Figures 2E to 2G). Notably, CMTM6-deficient B cells showed significantly reduced promotion of T cell-specific activation and proliferation (Figures 2E to 2G; Figure S6). Thus, we postulated that deficiencies in B-cell CMTM6 result in compromised T/B-cell interactions.

### CMTM6 affects B cell differentiation

Our another emphasis was on the impact of CMTM6 on B cell differentiation. We used flow cytometry to compare the percentages of each subpopulation of splenic B cells in WT and *Cmtm6* KO mice (Figure S7). We discovered that CMTM6 deficiencies had no significant effect on the proportion of total splenic B cells, as well as the percentages of B1 cells, B2 cells, marginal zone B cells, follicular B cells, immature B cells, mature B cells, and plasma cells (Figures S8A-D).

Notably, CMTM6-deficient mice showed considerably decreased percentages of splenic non-class-switching memory B cells, class-switching memory B cells, and germinal center (GC) B cells (Figures 2H and 2I). In particular, the percentage of GC B cells dropped by 69.01%. In mouse lymph nodes, the percentage of lymph node memory B cells and GC B cells was also much lower in CMTM6-deficient mice (Figures 2J and 2K). Moreover, by comparing splenic B cell subpopulations in *Cd274* KO mice and *Cmtm6*/*Cd274* double-KO mice, it was found that, in the absence of the PD-L1 axis, CMTM6 defects still resulted in a reduction of memory B cells and GC B cells (Figures 2L and 2M). These findings demonstrate that CMTM6 deficiency impairs B cell differentiation or survival to memory B cells and GC B cells, independently of PD-L1. CMTM6 may have impacted critical molecules in this biological process. Since mice do not express CD58, we left aside the question of whether CD58 plays a role.

### CMTM6 affects CD40-mediated B cell activation

To investigate the effect of CMTM6 on B cell immune function, we used five commonly used B cell agonists to compare the activation status of wild-type B cells and CMTM6-deficient B cells: the TLR4 agonist LPS, the TLR7 agonist Imiquimod, the TLR9 agonist Agatolimod, the BCR agonist anti-IgM antibody, and the CD40 agonist FGK45 antibody. Statistical analysis showed that B cell activation levels were similar between WT and *Cmtm6* KO splenocytes following TLR4, TLR7, TLR9, and BCR activation (Figures S9A-D), but differed considerably after anti-CD40 stimulation (Figures 3A-C). Specifically, B cells in the CMTM6-deficient splenocytes expressed considerably lower amounts of CD69, CD86, and major histocompatibility complex class II (MHC II) than those in the WT group, indicating decreased activation (Figures 3A-C).

**Figure 3.**
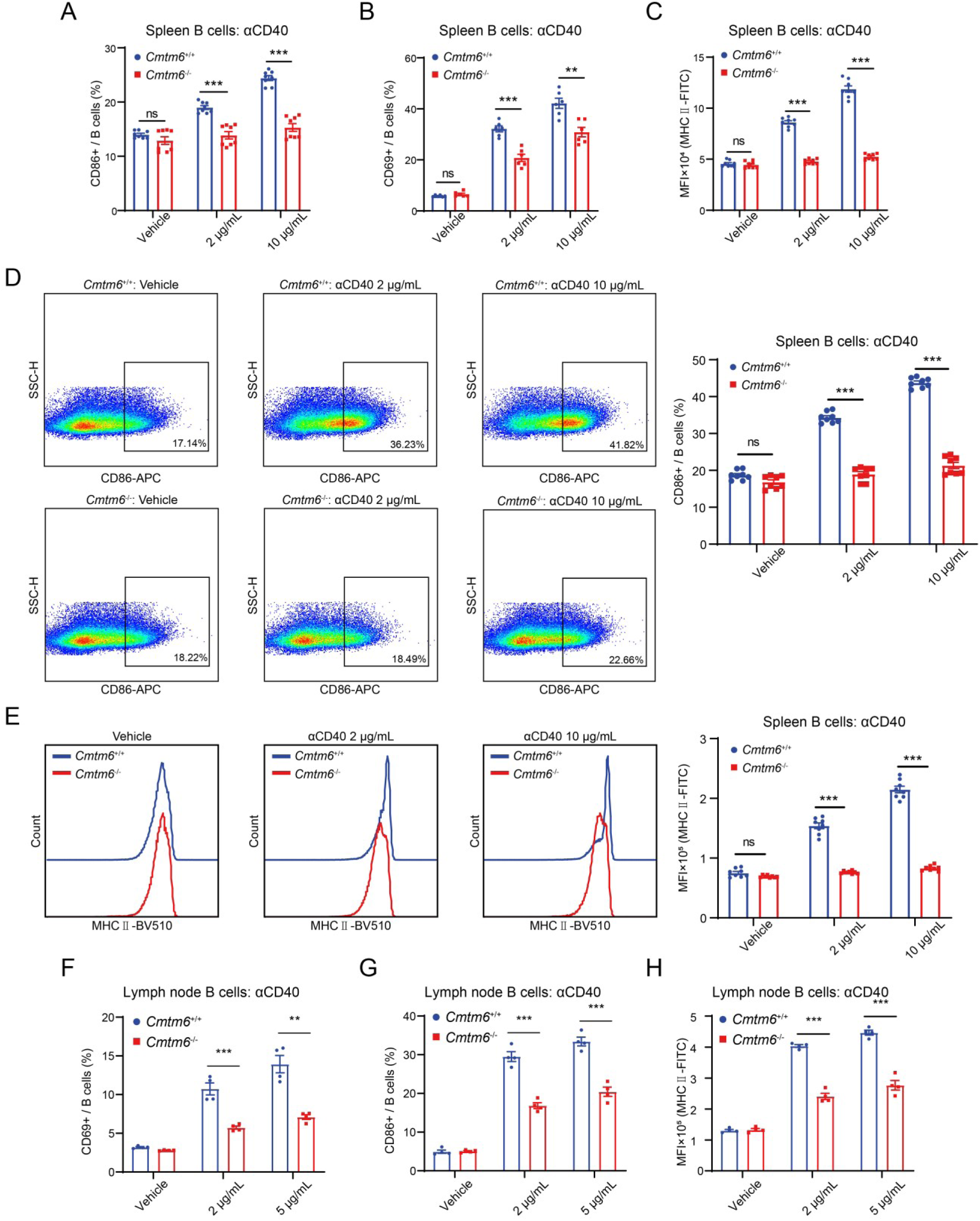
CMTM6 affects CD40-mediated B cell activation. (A-C) Statistical analysis of the proportion of CD86^+^ (A), the proportion of CD69^+^ (B), and the mean fluorescence intensity (MFI) of MHC II (C) in splenic B cells from WT or *Cmtm6* KO mice after the action of CD40 agonist (n = 8). (D and E) Activation analysis of sorted splenic B cells from WT or *Cmtm6* KO mice under CD40 agonism. Representative CD86^+^ population gating strategy and statistical analysis (D; n = 8). Representative histograms and statistical analysis of MHC II expression in B cells (E; n = 8). (F-H) Statistical analysis of the proportion of CD69^+^ (F), the proportion of CD86^+^ (G), and the MFI of MHC II (H) in lymph node B cells from WT or *Cmtm6* KO mice after the action of CD40 agonist (n = 4). The data are presented as the mean ± SEM. ** p < 0.01; *** p < 0.001; ns not significant by two-way ANOVA followed by Tukey’s multiple comparisons test.

To exclude out the interference of other immune cells in the splenocytes, we analyzed the activation of sorted B cells. We discovered that CMTM6-deficient B cells also exhibited significantly lower CD86 and MHC II expression compared to WT B cells when exposed to CD40 agonists (Figure 3D and 3E). Interestingly, there was no discernible increase in activation, even when compared to vehicle controls (Figure 3D and 3E). Clustering of CMTM6-deficient B was also notably decreased after FGK45 action (Figure S10). In addition, we arrived at the same conclusion after comparing the variations in lymph node B cells between WT and *Cmtm6* KO mice subsequent to anti-CD40 antibody activation (Figures 3F-H). These findings imply that CMTM6 does have an effect on B cell immunological function, and that the CD40 pathway may be a possible mechanism for CMTM6 regulating B cell immunity, such as T/B cell interactions, B cell differentiation and activation.

### CMTM6 maintains membrane levels of CD40 in B cells

Given the studies that have shown that CMTM6 interacts with two single-transmembrane proteins, PD-L1 and CD58, we speculated that CMTM6 may also interact with CD40, which is also a single-transmembrane receptor. Subsequently, by mining publicly available protein interactions data, we found that potential interactions between CMTM6 and CD40 were seen in yeast two-hybrid libraries, affinity-purified proteomics, and membrane proteomics of CMTM6 knockouts (Figure S11A). In addition, in open-source CRISPR screening data, we also found that KO of CMTM6 in human B-cell lymphoma cells Daudi affected the membrane level of CD40 (Figure S11B).

Therefore, we postulated that CMTM6 may also affect membrane levels of CD40 in B cells. Our findings indicate that, in comparison to WT mice, CMTM6 deficiency considerably decreased membrane CD40 in total splenic B cells and significantly decreased membranal CD40 in immature, mature, naïve, memory, and germinal center B cells (Figures 4A and 4B). In mouse lymph nodes, CMTM6 deficiencies also caused downregulation of CD40 membrane levels in broad B cell subsets (Figure 4C). We compared B-cell CD40 expression in *Cmtm6*/*Cd274* double-KO mice and *Cd274* KO mice to confirm the function of the PD-L1 axis in regulating CD40 membrane levels by CMTM6 (Figure 4D). Our findings revealed that CMTM6 defects similarly decreased membranal CD40 in the absence of the PD-L1 axis (Figure 4D). Besides, ablation of CMTM6 also reduced CD40 membrane levels in splenic B cells in MC38-OVA or B16-OVA tumor-bearing mice (Figure 4E). In addition, we also knocked down CMTM6 in human peripheral blood mononuclear cells (PBMCs) B cells by siRNA based on electrotransfection and similarly found that reduced expression of CMTM6 decreased CD40 levels in B cells (Figure S12). These findings show that CMTM6 is a regulatory molecule for B cell CD40 independently of PD-L1.

**Figure 4.**
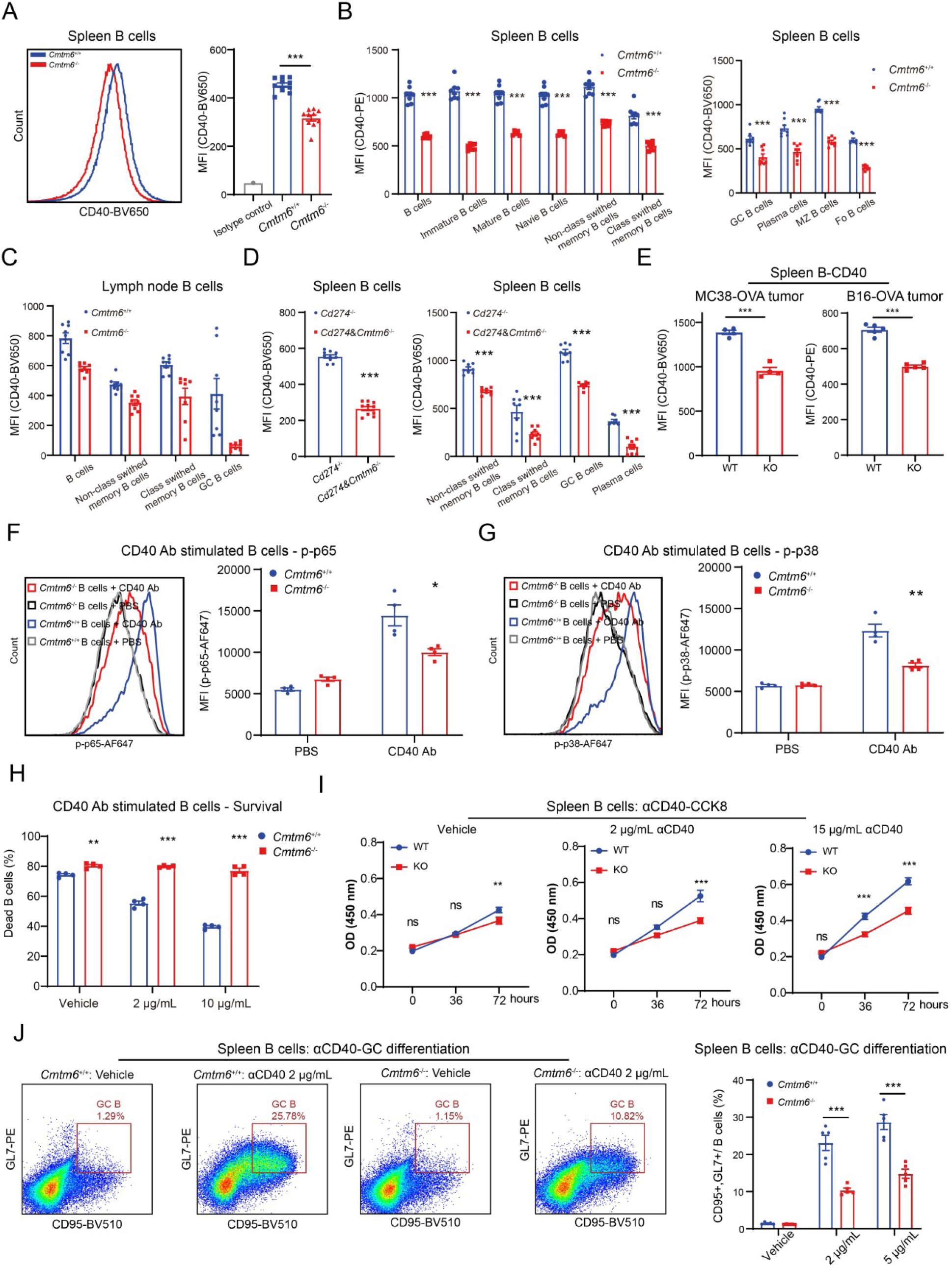
CMTM6 maintains membrane levels and function of CD40 in B cells. (A) Analysis of CD40 membrane levels in splenic B cells from WT or *Cmtm6* KO mice (n = 10). (B) Analysis of CD40 membrane levels in various subpopulations of splenic B cells from WT or *Cmtm6* KO mice by flow cytometry (n = 8). (C) Analysis of CD40 membrane levels in various subpopulations of lymph node B cells from WT or *Cmtm6* KO mice by flow cytometry (n = 8). (D) Analysis of CD40 membrane levels in various subpopulations of splenic B cells from *Cmtm6*/*Cd274* double-KO mice or *Cd274* KO by flow cytometry (n = 8). (E) Analysis of CD40 membrane levels in splenic B cells from WT or *Cmtm6* KO mice bearing MC38-OVA tumor (n = 4) or B16-OVA tumor (n = 5). (F and G) Analysis of phosphorylated p65 and p38 levels in splenic B cells from WT or *Cmtm6* KO mice after CD40 agonism by flow cytometry (n = 4). (H) Analysis of 7-AAD staining of dead splenic B cells from WT or *Cmtm6* KO mice after CD40 agonism for 72 h (n = 4). (I) Proliferation analysis of splenic B cells from WT or *Cmtm6* KO mice by CCK8 assay after CD40 agonism for 36 h or 72 h (n = 4). (J) Analysis of differentiation levels of germinal center B cells in splenic B cells from WT or *Cmtm6* KO mice after CD40 agonism for 72 h (n = 5). The data are presented as the mean ± SEM. * p < 0.05; ** p < 0.01; *** p < 0.001; ns not significant by unpaired t test or one-way/two-way ANOVA followed by Tukey’s multiple comparisons test.

### CMTM6 maintains function of CD40 in B cells

We therefore postulated that CMTM6 may influence CD40 downstream signaling and function through regulating its membrane levels. We sorted primary splenic B cells from WT and *Cmtm6* KO mice and acted on them with CD40 agonist for 15 min, respectively. Phospho flow cytometry revealed that CMTM6-deficient B cells had significantly reduced levels of phosphorylated p65 and p38, suggesting that CMTM6 indeed affects CD40 signaling in B cells (Figures 4F and 4G).

During in vitro culture, primary B cells quickly undergo an apoptotic process, and CD40 signaling can increase anti-apoptotic and proliferation of B cells. B cells from WT mice showed increased survival after 72 hours of CD40 agonist treatment due to a dose-dependent anti-apoptotic effect, which was not observed in the CMTM6 deficiency group (Figure 4H). CD40 agonist-induced B cell proliferation was also strongly influenced by CMTM6 KO in CCK8 assays (Figure 4I). CD40 signaling can enhance the development of GC B cells. We observed a dose-dependent differentiation of WT B cells into GL7^+^ CD95^+^ GC B cells when treating primary B cells in vitro with CD40 agonists (Figure 4J). However, the differentiation of B cells was notably decreased in CMTM6 KO mice (Figure 4J).

Collectively, CMTM6 affects CD40 signaling-mediated B cell activation, proliferation, survival, and differentiation through its effect on B cell CD40 membrane levels.

### CMTM6 depends on CD40 signaling for modulating B cell function

To further validate the role of CD40 signaling in CMTM6-regulated B-cell immunity, we observed the effects of CD40L-blocking antibody on CMTM6-deficient B-cell effects. In the T/B cell co-culture model, blockade of the CD40/CD40L interaction resulted in a substantial attenuation of the promotion of OT-II CD4+ T cell activation and proliferation by B cells loaded with OVA (Figure 5A and 5B). Discrepancies between the CD19-iCre B cell group and the BKO B cell group continued to be observed after anti-CD40L treatment (Figure 5A and 5B). However, these results at least suggest that CD40 signaling plays an important role in T/B interactions and that CMTM6 can be implicated in T/B interactions by modulating CD40 signaling.

**Figure 5.**
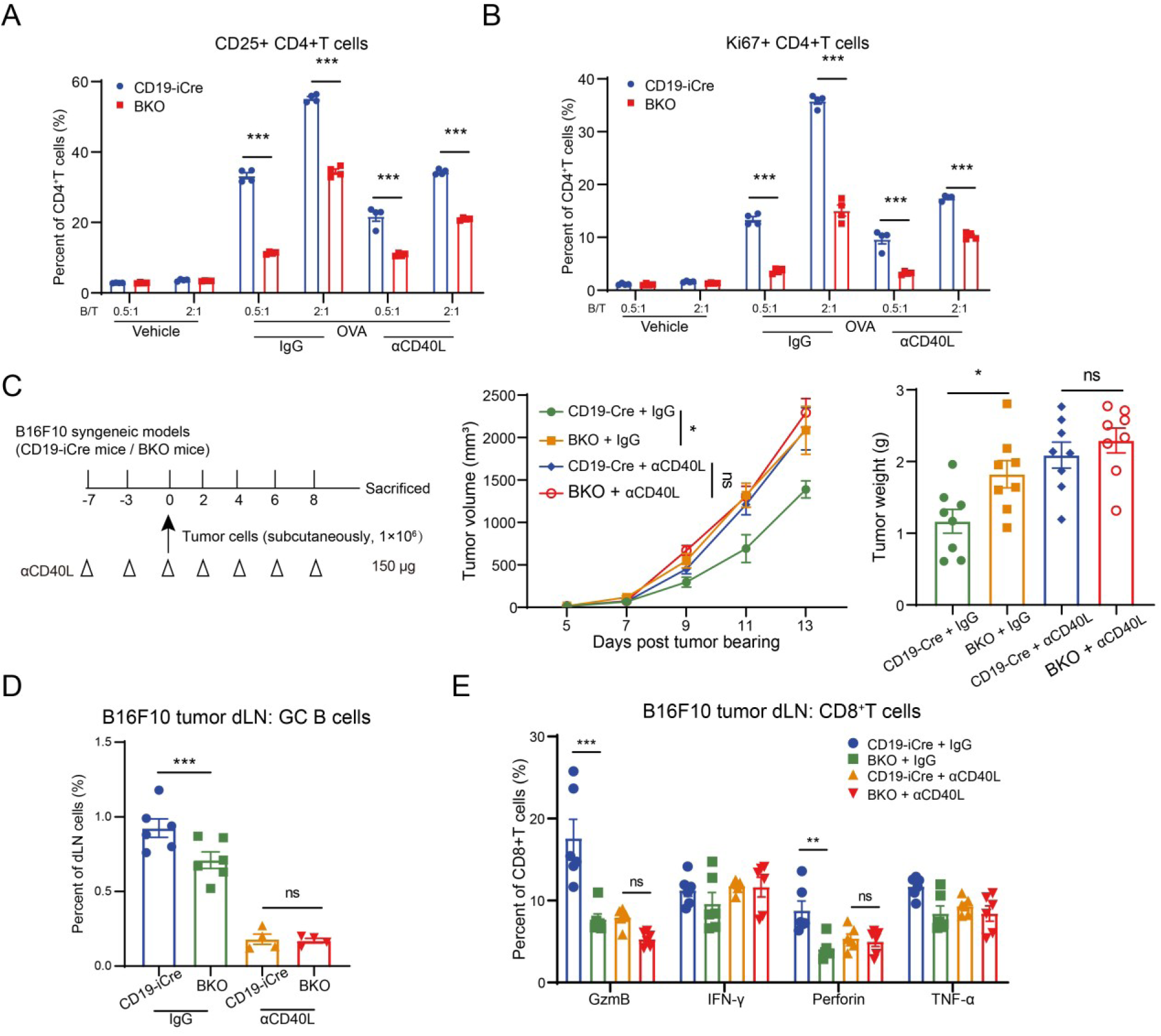
CMTM6 is dependent on CD40 signaling to regulate B cell function. (A and B) The activation levels (A: CD25^+^) and the proliferation levels (B: Ki67^+^) of OT-Ⅱ CD4^+^ T cells with IgG or anti-CD40L antibody treatment when co-cultured with CD19-iCre or BKO B cells (n = 4). (C) Tumor growth and tumor weight of B16F10 tumors in CD19-iCre or BKO mice with IgG or anti-CD40L antibody treatment (n = 8). (D) The percentage of GC B cells in dLNs of B16F10 tumor-bearing CD19-iCre or BKO mice with IgG or anti-CD40L antibody treatment (n = 6). (E) Effector molecule levels of dLN CD8^+^ T cells of B16F10 tumor-bearing CD19-iCre or BKO mice with IgG or anti-CD40L antibody treatment (n = 6). The data are presented as the mean ± SEM. * p < 0.05; ** p < 0.01; *** p < 0.001; ns not significant by one-way/two-way ANOVA followed by Tukey’s multiple comparisons test.

In vivo, CD40 signaling suppression led to both an acceleration of tumor progression and the elimination of tumor growth and weight disparities between the CD19-iCre and BKO groups (Figure 5C). Further, we analyzed the GC B-cell ratio and CD8^+^ T-cell effect in the dLN. Anti-CD40L administration significantly reduced the level of GC B-cells and eliminated the difference between the CD19-iCre group and the BKO group (Figure 5D). Similarly, the levels of effector molecules such as IFN-γ produced by CD8^+^ T were no longer affected by CMTM6 knockout in B cells after CD40/CD40L blockade (Figure 5E). Taken together, CMTM6 is at least partially dependent on CD40 signaling to regulate T/B cell interactions, GC B cell generation, and B cell antitumor effects.

### CMTM6 is an interacting molecule of CD40

The foregoing findings suggest that CMTM6 is a new regulatory molecule for B cell immunity, and that the CMTM6-CD40 axis controls B cell function. Therefore, we aim to investigate the mode of CMTM6-CD40 interaction in depth. First, at the transcriptional level, *Cmtm6* KO did not affect *Cd40* expression in mouse spleen B cells (Figure 6A). We then pulled CD40 protein based on the anti-CMTM6 antibody in mouse spleen cell samples or based on the anti-Flag antibody in HEK293T cells with CMTM6/CD40 co-overexpression by Co-IP assay, and the results showed that CMTM6 and CD40 did have direct interaction (Figure 6B).

**Figure 6.**
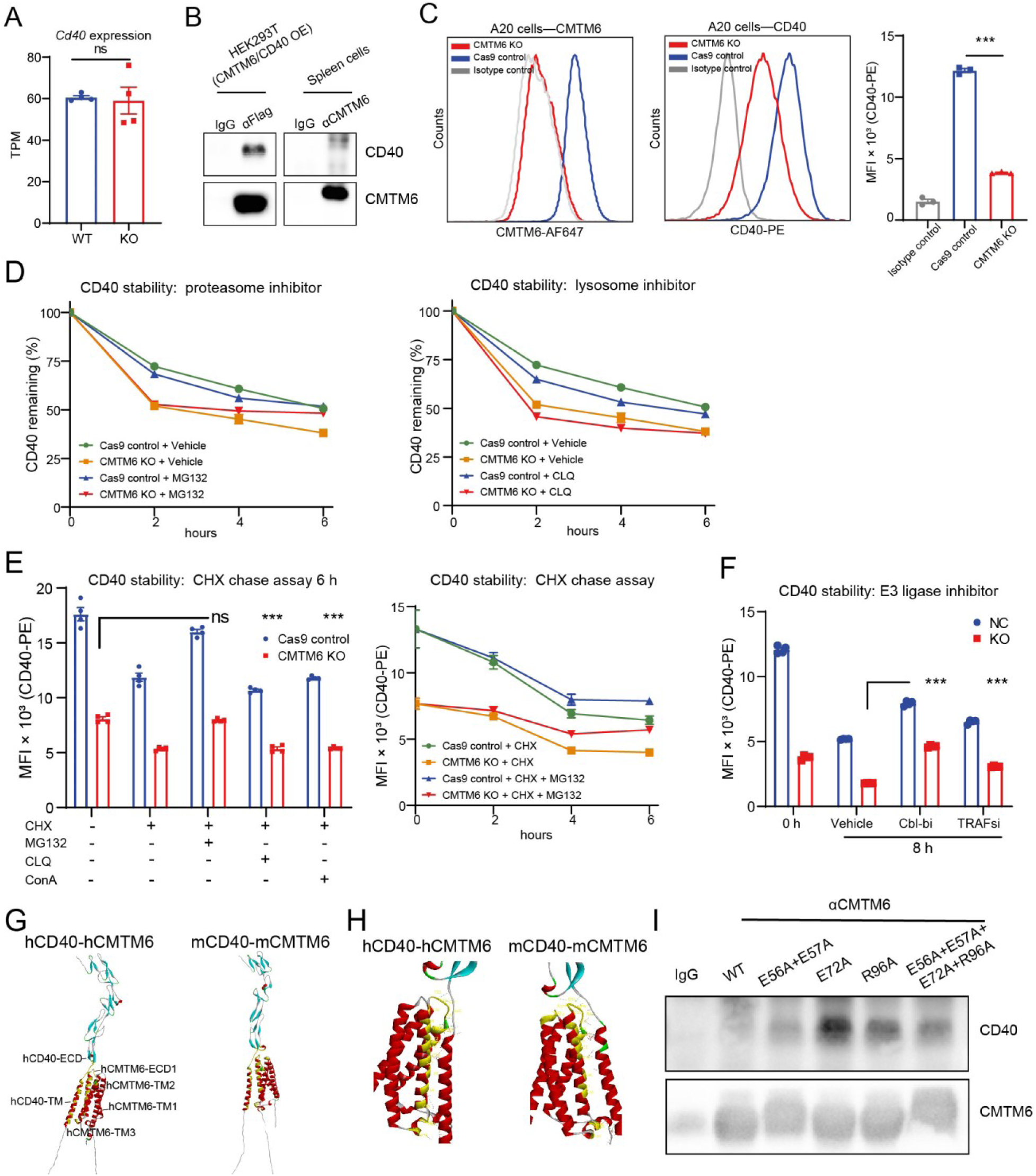
CMTM6 interacts with CD40 and inhibits its degradation by the ubiquitin-proteasome system. (A) Transcripts per million (TPM) values of *Cd40* in mouse spleen B cells by RNA-seq (n = 4). (B) Co-IP analysis of CMTM6/CD40 protein in mouse splenocyte samples and HEK293T cells with CMTM6/CD40 overexpression. (C) Analysis of CD40 membrane levels in Cas9 control A20 cells or CMTM6 KO A20 cells by flow cytometry (n = 3). (D) Analysis of CD40 membrane levels by flow cytometry in Cas9 control A20 cells or CMTM6 KO A20 cells pre-stained for CD40 and treated with MG132 or CLQ (n = 4). (E) Analysis of CD40 membrane levels by flow cytometry in Cas9 control A20 cells or CMTM6 KO A20 cells and treated with CHX, MG132, CLQ or ConA (n = 4). (F) Analysis of CD40 membrane levels by flow cytometry in Cas9 control A20 cells or CMTM6 KO A20 cells pre-stained for CD40 and treated with Cbl-b inhibitor Cbl-b-IN-5 or TRAFs inhibitor BC-1215 (n = 4). (G) AlphaFold2-predicted structures of full-length complexes of human or mouse CMTM6/CD40. (H) Key interaction sites predicted by AlphaFold2 for human or mouse CMTM6/CD40. (I) Co-IP analysis of CMTM6/CD40 protein in HEK293T cells with CD40 and different CMTM6 mutants overexpression. The data are presented as the mean ± SEM. *** p < 0.001; ns not significant by unpaired t test or one-way/two-way ANOVA followed by Tukey’s multiple comparisons test.

To further investigate the mechanism by which CMTM6 regulates CD40 stability, we constructed the mouse B lymphoma cell line A20 with Cmtm6 KO and again verified by flow cytometry that defects in CMTM6 affect CD40 membrane levels (Figure 6C). In A20 cells, by membrane CD40 degradation assays, we found that proteasome inhibition by MG132 significantly slowed down the rapid CD40 degradation caused by *Cmtm6* KO, whereas lysosomal inhibition by chloroquine did not (Figure 6D). A similar phenomenon was observed in Cycloheximide (CHX) chase assays, suggesting that CMTM6 regulates CD40 stability mainly through the proteasome pathway (Figure 6E). Due to the scarcity of publications directly addressing CD40 degradation, we found papers indicating that Cbl-b and TRAFs are associated with the levels of the CD40 protein complex^23–26^, and that both molecules exhibit E3 ubiquitin ligase activity. We discovered that inhibiting Cbl-b and TRAFs activities might significantly impede the accelerated CD40 degradation induced by Cmtm6 knockout by the use of both inhibitors, suggesting that CMTM6 may block the ubiquitinated degradation of CD40 by E3 enzymes (Figure 6F).

AlphaFold2 was utilized to predict the structures of human and mouse CMTM6/CD40 complexes, human and mouse CMTM6/PD-L1 complexes, and human CMTM6/CD58 complexes (Figures 6G and S13). The interaction interfaces within these complexes were also analyzed. The sites of interaction in CMTM6 were highly similar between CD40/CD58/PD-L1, and the main regions of interaction were located in the first transmembrane domain, the second transmembrane domain, and the first extracellular domain of the CMTM6 protein (Figure 6H). The sites of interaction on CD40 were mainly located in the transmembrane domain and the proximal transmembrane region. By alanine mutation scanning, we found that in CMTM6, sites such as E56, E57, E72, and R96 have a strong influence on the CMTM6/CD40 interaction, and mutations can enhance the binding of the two. Subsequently, we constructed four CMTM6 mutants, and after transfecting HEK293T cells, we found by Co-IP that mutations at these sites did enhance CMTM6/CD40 interactions, suggesting that CMTM6 most likely binds to CD40 via TM1, ECD1, TM2, as predicted by AlphaFold (Figure 6I).

### B-cell CMTM6 impacts the antitumor efficacy of CD40 agonists

The above results illustrate that CMTM6 affects B-cell immunity by interacting with CD40 and maintaining B-cell CD40 membrane levels and regulating CD40 signaling and function. In the B16F10 subcutaneous tumor model, we discovered that CD20 antibody-mediated deletion of B cells in mice significantly affected the antitumor activity of CD40 agonists (Figure 7A), and similar phenomena were observed in the CT26 and MC38 models (Figures 7B and 7C), implying that B cells are one of the pharmacological target cells for CD40 agonists. Next, in the B16F10 model and MC38 model, we found that KO of host CMTM6 significantly reduced the in vivo antitumor activity of CD40 agonists (Figures 7D and 7E). Considering our findings that CMTM6 regulates B-cell CD40 and B-cell immunity, we hypothesized that CMTM6 may influence the antitumor efficacy of CD40 agonists by regulating CD40 signaling in B cells. Of course, we also verified the hypothesis in BKO mice. The efficacy of the CD40 agonist was highly diminished by B-cell CMTM6 deficiency, and the IR for tumor weight was reduced from 88.84% to 67.17% (Figures 7F). Meanwhile, considering the influence of B-cell CMTM6 on the antitumor effect of T cells, we also evaluated the efficacy of a PD-1 inhibitor in BKO mice, and the results showed that the tumor suppression effect of PD-1 blockade was also significantly inhibited after B-cell CMTM6 deficiency (Figures 7G). These results suggest a correlation between B-cell CMTM6 and the antitumor efficacy of CD40 agonist and ICB therapies, which could be validated in data from clinical tumor samples in further studies.

**Figure 7.**
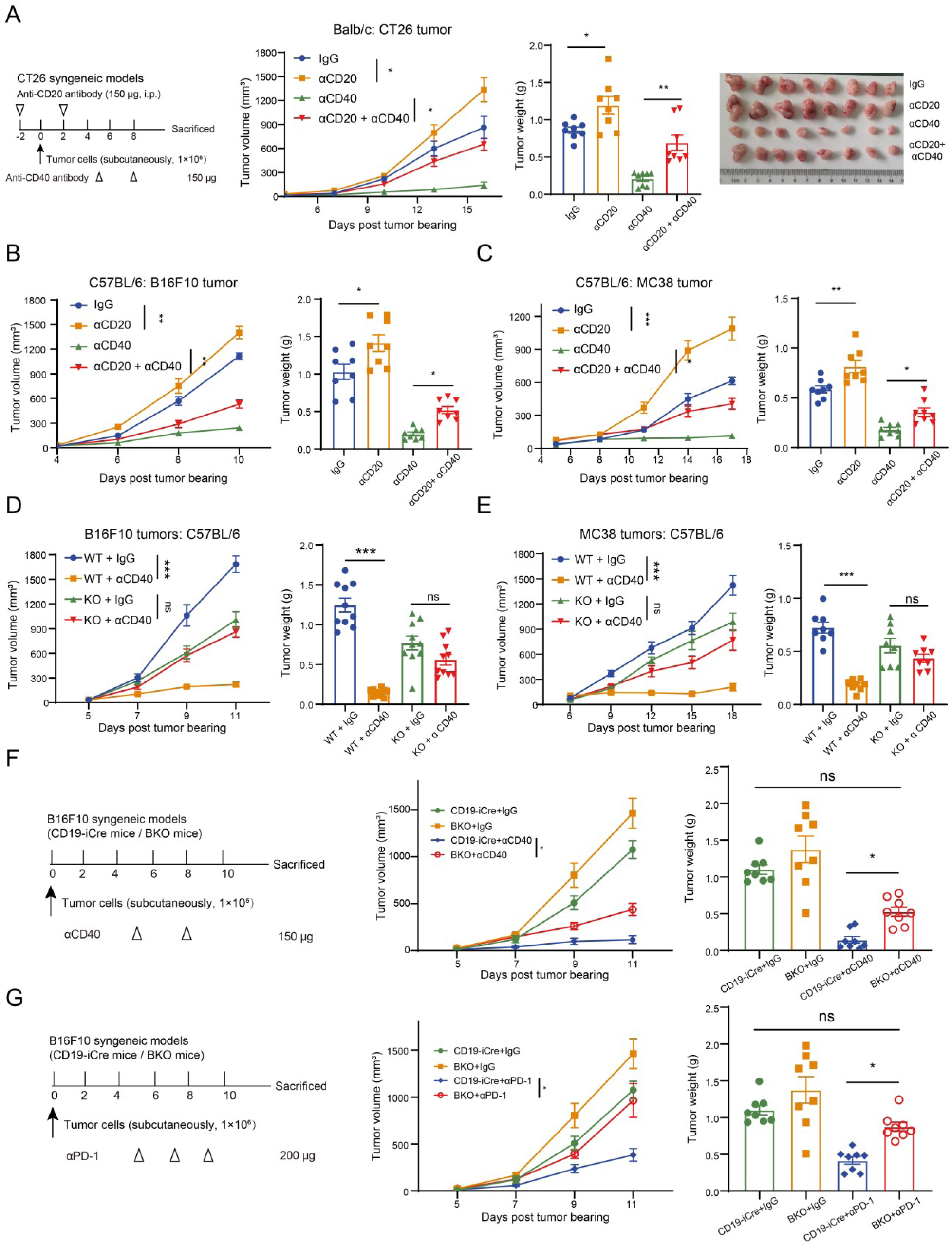
B-cell CMTM6 impacts the antitumor efficacy of CD40 agonists. (A) Effect of B-cell deletion on the efficacy of CD40 agonists against CT26 tumors (n = 8). (B) Effect of B-cell deletion on the efficacy of CD40 agonists against B16F10 tumors (n = 8). (C) Effect of B-cell deletion on the efficacy of CD40 agonists against MC38 tumors (n = 8). (D) Comparison of CD40 agonist antitumor efficacy in WT and *Cmtm6* KO mice in the B16F10 tumor model (n = 10). (E) Comparison of CD40 agonist antitumor efficacy in WT and *Cmtm6* KO mice in the MC38 tumor model (n = 8). (F) Tumor growth and tumor weight of B16F10 tumors in CD19-iCre or BKO mice with IgG or anti-CD40 antibody treatment (n = 8). (G) Tumor growth and tumor weight of B16F10 tumors in CD19-iCre or BKO mice with IgG or anti-PD-1 antibody treatment (n = 8). The data are presented as the mean ± SEM. * p < 0.05; ** p < 0.01; *** p < 0.001; ns not significant by one-way ANOVA followed by Tukey’s multiple comparisons test.

## DISCUSSION

In this work, we investigated the influence of the CMTM6-CD40 axis in B cell differentiation, B cell immunity and B cell anti-tumor immune effects.

In previous studies, B cells have often been considered to be bystanders to tumor immunity or to have pro-tumorigenic effects. Initially, researchers discovered that the anti-tumor effects of T cells were boosted in the B-cell-deficient mice μMT^27,28^. The researchers also identified a subpopulation of B cells with immunosuppressive functions called regulatory B cells (Bregs)^29^. Bregs, which expresses IL-10 as a marker, can suppress anti-tumor immunity by secreting IL-10, IL-35, TGF-β, IL-1β, or expressing checkpoint molecules such as PD-1 and PD-L1^30–32^. In addition, IgA antibodies produced by B cells have also been reported to dampen the cytotoxicity of CD8^+^ T cells^33^. But as we mentioned earlier, the antitumor role of B cells has received more attention in recent years. Moreover, the latest researches have raised questions about the role of tumor-promoting factors produced by B cells. The researchers found that conditional KO of IL-10 in B cells did not inhibit tumor growth^9^. IgA antibodies have also been reported to enter tumor cells via transcytosis, block oncogenic signals, and promote T cell-mediated cytotoxicity^34^. Our findings reaffirm that total B cells possess anti-tumor effects and can promote intratumor T cell infiltration and effects. We also discovered that CD40 agonists rely on B cells for effectiveness, indicating that B cells may play a crucial role in tumor immunotherapy. Of course, further investigation is required to understand the role and function of B lymphocytes in the tumor microenvironment in order to maximize their potential in tumor immunotherapy.

We identified CMTM6 as a novel regulatory molecule of B cell intrinsic CD40. CD40 signaling is crucial for B cell proliferation, survival, activation, differentiation, and antibody production^35^. By modulating CD40 signaling, CMTM6 was involved in B cell survival, activation, differentiation, proliferation, and antitumor immunity. In line with findings reported in previous research involving *Cd40* KO mice or *Cd40l* KO mice^36,37^, *Cmtm6* KO mice displayed atypical quantities of germinal center B cells and memory B cells. In recent years, several studies have also validated that strong CD40/CD40L interactions are critical for B cell differentiation^38,39^.

Interestingly, although KO of CMTM6 resulted in only about a 30% decrease in CD40 membrane levels, CD40 signaling and function were significantly attenuated. Our hypothesis posits that CMTM6, apart from its role in CD40 membrane level regulation, might also participate in the interplay of signals between CD40L and CD40 as a component of the CD40 signaling complex.

Our results suggest that CMTM6 is a novel interacting molecule of CD40. Recent studies have identified CMTM6 as a molecule that interacts with numerous transmembrane proteins, including PD-L1, CD58, and HER2^16–19,40^. CMTM6 regulates the membrane levels of these proteins in the post-translational modification pathway, impacting tumor growth and tumor immunity. Considering the predicted structure of CMTM6 and the similar interaction patterns of CMTM6 with other proteins, we hypothesize that CMTM6 might serve as a chaperone protein responsible for sustaining the stability of various transmembrane proteins. Interestingly, molecules regulated by CMTM6 exhibit variable functions in the progression of tumors, indicating the intricate function of CMTM6 in the tumor microenvironment.

In summary, our work demonstrates that B-cell intrinsic CMTM6 regulates CD40 signaling and function via maintaining the B-cell membranal CD40 through a post-translational modification pathway, thereby affecting B-cell differentiation and function, anti-tumor B-cell immunity, and the efficacy of CD40 agonists. These studies will broaden the knowledge of the anti-tumor effects of B cells and the regulatory mechanism for CD40, elucidate the role and mechanism of CMTM6 in the TME, and inspire the value of B cell CMTM6 for potential clinical applications.

### Limitations of the study

In the present study, we did not validate the regulatory effect of CMTM6 on B-cell antitumor immunity in an in-situ tumor model and an immune-humanized tumor model due to the limitation of conditions. Besides, considering that CMTM6 and CD40 are also highly expressed in macrophages, neutrophils, etc., we should also explore the role of the CMTM6-CD40 axis in other immune cells in future studies.

## SUPPLEMENTAL INFORMATION

Figures. S1 to S13

## Supporting information

Main text

## ACKNOWLEDGMENTS

We would like to thank Professor Min Huang from the Shanghai Institute of Materia Medica, Chinese Academy of Sciences, who provided scientific advices. This work was supported by the China Postdoctoral Science Foundation (2023M743659 and 2023TQ0364), the Shanghai Post-doctoral Excellence Program (2023699), the Postdoctoral Fellowship Program of CPSF (GZB20230797), Shanghai Rising-Star/Sailing Program (24YF2756000) and the National Natural Science Foundation of China (32400749). All the schematics are were created with BioRender.com.

## AUTHOR CONTRIBUTIONS

Conceptualization, L-K.G. and Y.L.; Methodology, Y.L. and R.C.; Formal Analysis, Y.L.; Investigation, Y.L., R.C, W.C., F.L. and L-H.G.; Resources, L-K.G., J.C., J.S., Y.L.; Writing – Original Draft, Y.L.; Writing – Review & Editing, L-K.G. and Y.L.; Supervision, L-K.G.; Funding Acquisition, L-K.G. and Y.L.

## STAR*METHODS

### KEY RESOURCES TABLE

| REAGENT or RESOURCE | SOURCE | IDENTIFIER |
| --- | --- | --- |
| **Antibodies** |  |  |
| Ms CD45R/B220 PerCP-Cy5.5 RA3-6B2 | BD Bioscience | Cat# 552771 |
| BV650 Rat Anti-Mouse CD45R/B220 (RA3-6B2) | BD Bioscience | Cat# 563893 |
| PE Hamster Anti-Mouse CD27 (LG.3A10) | BD Bioscience | Cat# 558754 |
| PE-Cy7 Hamster Anti-Mouse CD27 (LG.3A10) | BD Bioscience | Cat# 563604 |
| Ms IgM BV605 II/41 | BD Bioscience | Cat# 743325 |
| APC Rat Anti-Mouse IgD (11-26c.2a) | BD Bioscience | Cat# 560868 |
| FITC Rat Anti-Mouse CD19 (1D3) | BD Bioscience | Cat# 553785 |
| PE-Cy7 Rat Anti-Mouse CD43 (S7) | BD Bioscience | Cat# 562866 |
| Ms CD5 BV510 53-7.3 | BD Bioscience | Cat# 563069 |
| MS CD38 BV786 90/CD38 | BD Bioscience | Cat# 740887 |
| BV421 Rat Anti-Mouse CD24 (M1/69) | BD Bioscience | Cat# 562563 |
| Ms CD40 BV650 3/23 | BD Bioscience | Cat# 740492 |
| PE Rat Anti-Mouse T- and B-Cell Activation Antigen (GL7) | BD Bioscience | Cat# 561530 |
| BV786 Rat Anti-Mouse CD23 (B3B4) | BD Bioscience | Cat# 563988 |
| BV421 Rat Anti-Mouse CD21/CD35 (7G6) | BD Bioscience | Cat# 562756 |
| PE Rat Anti-Mouse CD40(3/23) | BD Bioscience | Cat# 561846 |
| BV421 Mouse Anti-Human CD40(5C3) | BD Bioscience | Cat# 563396 |
| Brilliant Violet 650 anti-human CD19 | Biolegend | Cat# 302238 |
| Ms CD4 BV421 GK1.5 | BD Bioscience | Cat# 562891 |
| Ms CD8a BV510 53-6.7 | BD Bioscience | Cat# 563068 |
| Ms CD3e APC-Cy7 145-2C11 | BD Bioscience | Cat# 557596 |
| FITC Rat Anti-Mouse CD45 (30-F11) | BD Bioscience | Cat# 553079 |
| Ms CD86 APC GL1 | BD Bioscience | Cat# 558703 |
| PE Hamster Anti-Mouse CD69 (H1.2F3) | BD Bioscience | Cat# 561932 |
| Ms TNF BV650 MP6-XT22 | BD Bioscience | Cat# 563943 |
| BV786 Rat Anti-Mouse IFN-γ XMG1.2 | BD Bioscience | Cat# 563773 |
| Ms I-A/I-E BV510 2G9 | BD Bioscience | Cat# 743871 |
| BV650 Rat Anti-Mouse CD138 (281-2) | BD Bioscience | Cat# 564068 |
| BV510 Hamster Anti-Mouse CD95 (Jo2) | BD Bioscience | Cat# 563646 |
| APC Rat anti-Mouse CD138 (281-2) | BD Bioscience | Cat# 558626 |
| ANTI-MOUSE GRANZYME B (NGZB) PE-CYANINE | eBioscience | Cat# 25-8898-82 |
| ANTI-MO PERFORIN (EBIOOMAK-D) APC | eBioscience | Cat# 17-9392-80 |
| Anti-CMTM6 antibody [EPR23015-45] | Abcam | Cat# ab264067 |
| Goat Anti-Rabbit IgG H&L (Alexa Fluor® 647) | Abcam | Cat# ab150079 |
| Goat Anti-Mouse IgM, F(ab')<sub>2</sub> Antibody | CST | Cat# 46315S |
| Anti-CD40 antibody [EPR18005-35] | Abcam | Cat# ab188181 |
| DYKDDDDK Tag (FLAG) Antibody | MCE | Cat# HY-P80111 |
| DDDDK-Tag Rabbit mAb | ABclonal | Cat# AE092 |
| InVivoMAb anti-mouse CD20 | BioXcell | Cat# BE0356 |
| InVivoMAb anti-mouse CD40 | BioXcell | Cat# BE0016 |
| InVivoMAb anti-mouse CD40L (CD154) | BioXcell | Cat# BE0017 |
| InVivoMAb polyclonal Armenian hamster IgG | BioXcell | Cat# BE0091 |
| **Bacterial and virus strains** |  |  |
| DH5α | Vazyme | Cat# C502-02 |
| Stbl3 | Absin | Cat# abs60045 |
| **Chemicals, peptides, and recombinant proteins** |  |  |
| Imiquimod | MCE | Cat# HY-B0180 |
| Brefeldin A (BFA) Solution | BD Pharmingen | Cat# 347688 |
| MG132 | MCE | Cat# HY-13259 |
| Chloroquine phosphate (CLQ) | MCE | Cat# HY-17589 |
| Concanamycin A (ConA) | MCE | Cat# HY-N1724 |
| Cycloheximide (CHX) | MCE | Cat# HY-12320 |
| Cbl-b-IN-5 | MCE | Cat# HY-155120 |
| BC-1215 | MCE | Cat# HY-117937 |
| Mouse IL-2 | Genscript | Cat# Z02764 |
| Mouse IL-21 | MCE | Cat# HY-P700199AF |
| Mouse BAFF | MCE | Cat# HY-P700161AF |
| Mouse IL-4 | MCE | Cat# HY-P70653 |
| LPS (055:B5) | Merck | Cat# L6529 |
| Agatolimod | MCE | Cat# HY-150218 |
| 7-AAD | BD Pharmingen | Cat# 559925 |
| Fixable Viability Stain 575V (FVS575V) | BD Horizon | Cat# 565694 |
| **Critical commercial assays** |  |  |
| Cell Counting Kit (CCK-8) | Yeasen | Cat# 40203ES60 |
| Cytofix/Cytoperm Soln Kit | BD Pharmingen | Cat# 554714 |
| Transcription Factor Buffer Set | BD Pharmingen | Cat# 562574 |
| Phosflow™ Perm Buffer III | BD Pharmingen | Cat# 558050 |
| Cytofix™ Fixation Buffer | BD Pharmingen | Cat# 554655 |
| EasySep™ Mouse Pan-B Cell Isolation Kit | Stemcell | Cat# 19844 |
| EasySep™ Mouse CD4<sup>+</sup> T Cell Isolation Kit | Stemcell | Cat# 19852 |
| Red Blood Cell Lysis Buffer | Yeasen | Cat# 40401ES60 |
| **Experimental models: cell lines** |  |  |
| HEK293T | Procell | Cat# CL-0005 |
| CT26 | Procell | Cat# CL-0071 |
| B16F10 | Procell | Cat# CL-0319 |
| MC38 | BNCC | Cat#<br>BNCC337716 |
| B16-OVA | This lab | N/A |
| MC38-OVA | This lab | N/A |
| **Experimental models: organisms/strains** |  |  |
| Mouse: BALB/c | Shanghai Sipp-BK | N/A |
| Mouse: C57BL/6 | Shanghai Sipp-BK | N/A |
| Mouse: *Cmtm6* KO | GemPharmatech | T013922 |
| Mouse: *Cd274* KO | GemPharmatech | T009724 |
| Mouse: *Cd274&Cmtm6* KO | This lab | N/A |
| Mouse: *Cd19*-iCre | GemPharmatech | T006948 |
| Mouse: *Cmtm6* flox | GemPharmatech | T013033 |
| Mouse: OT-II | the Jackson<br>Laboratory | 004194 |
| **Software and algorithms** |  |  |
| Graphpad Prism 8.0 software | GraphPad Software, Inc | http://www.graphpad.com/scientificsoftware/prism/ |
| NovoExpress 1.6.0 | Agilent | https://www.agilent.com.cn/zh-cn/product/research-flow-cytometry/flow-cytometry-software |
| Adobe Illustrator 2017 | Adobe | https://www.adobe.com/adobe/illustrator |
| Discovery studio 2016 | BIOVIA | https://www.3ds.com/products/biovia/discovery-studio |
| AlphaFold2-multimer | ColabFold | https://colab.research.google.com/github/sokrypton/ColabFold/blob/main/AlphaFold2.ipynb |

## RESOURCE AVAILABILITY

### Lead contact

Further information and requests for materials should be directed to the Lead Contact: Likun Gong.

### Materials availability

This study did not generate new unique reagents.

### Data and code availability

This study did not generate new codes. Data are all available in the main text or the supplementary materials.

## EXPERIMENTAL MODEL AND SUBJECT DETAILS

### Cell lines

CT26, B16F10 and HEK293T cell lines were purchased from Procell. MC38 cell line was purchased from BNCC. B16-F10 and HEK293T cells were cultured in Dulbecco’s Modified Eagle Medium (DMEM; Meilunbio) supplemented with 1% penicillin/streptomycin (P/S; Invitrogen) and 10% heat-inactivated fetal bovine serum (FBS; Life Technologies). CT26 and MC38 were cultured in RPMI1640 medium (Meilunbio) containing 1% P/S and 10% FBS. MC38-OVA cell line and B16-OVA cell line were constructed by lentiviral systems. All cells were cultured at 37°C in a 5% CO_2_ humidified atmosphere.

### Animals

Female (Six-to eight-week-old) BALB/c and C57BL/6 mice were purchased from the Shanghai Slack Animal. *Cmtm6*^-/-^, *Cd274*^-/-^, *Cd19*-iCre and *Cmtm6*^flox^ mice were purchased from GemPharmatech. CMTM6 and PD-L1 double knockout mice were obtained by *Cmtm6*^-/-^ and *Cd274*^-/-^ mice hybridization. The *Cmtm6*^flox^ mice were bred to the *Cd19*-iCre mice to generate *Cmtm6*^fl/fl^ *Cd19*^cre+^ mice. OT-II transgenic mice were purchased from the Jackson Laboratory. All mice were maintained under specific pathogen-free (SPF) conditions in the animal facility of the Shanghai Institute of Materia Medica, Chinese Academy of Sciences (SIMM). Animal care and experiments were performed in accordance with the protocols approved by the Institutional Laboratory Animal Care and Use Committee (IACUC).

## METHOD DETAILS

### Primary B cell isolation and culture

WT or *Cmtm6*^-/-^ C57BL/6 mice were sacrificed, and the spleens or lymph nodes were collected for single-cell suspensions. Single-cell suspensions were obtained by grinding tissues through the rubber end of a 2 mL syringe (92311610, Sinopharm) and then were filtered through 70-μm Cell Strainers (WHB-70um, WHB). Single cells were subject to erythrocyte removal by red blood cell lysis buffer (40401ES60, Yeasen).

Highly purified B cells was isolated from splenocytes by immunomagnetic negative selection. The sorting process follows the instructions of the EasySep™ Mouse B Cell Isolation Kit (19854, Stemcell). Briefly, splenocytes were blocked by rat serum. Cells were added with isolation cocktail and incubated for 10 min at room temperature. Then, cells were added with RapidSpheres and incubated for 2.5 min at room temperature. Finally, B cells were sorted out by magnet and washed for subsequent experiments.

Primary B cells were cultured in RPMI1640 medium (Meilunbio) containing 1% P/S, 10% FBS, 50 ng/mL of IL-2 (Z02764, GenScript), 10 ng/mL of IL-4 (HY-P70653, MCE), 10 ng/mL of IL-21 (HY-P700199AF, MCE), and 10 ng/mL of BAFF (HY-P700161AF, MCE).

### Activation analysis of primary B cells

1×10^6^ of splenocytes or lymph node cells or primary B cells were added to 24-well plates and were stimulated with the TLR4 agonist LPS (L6529, Merck), the TLR7 agonist Imiquimod (HY-B0180, MCE), the TLR9 agonist Agatolimod (HY-150218, MCE), the BCR agonist anti-IgM antibody (46315S, CST), or the CD40 agonist FGK45 antibody (BE0016-2, BioXcell) for 24 h separately for activation phenotype analysis. Dissolution and preparation of agonists were performed following the manufacturer’s recommendations. Indicators of activation of primary B cells, CD69, CD86 and MHC II, were detected by flow cytometry.

### In vitro proliferation and survival analysis of primary B cells

For the cell proliferation assay, 1×10^5^ of primary B cells (100 μL) were seeded in 96-well plates. WT or Cmtm6 KO B cells were added with PBS, 2 μg/mL of CD40 agonist, or 15 μg/mL of CD40 agonist, respectively. 100 μL of CCK8 (40203ES60, Yeasen) was added at 0 h, 36 h, 72 h, respectively, and the signal was read at OD 450 nm by an automatic microplate reader SpectraMax (Molecular Devices).

For the survival assay, 1×10^6^ of WT or Cmtm6 KO primary B cells were added to 24-well plates and were stimulated with PBS, 2 μg/mL of CD40 agonist, or 10 μg/mL of CD40 agonist, respectively. After 72 h, B cells were collected and stained with 5 μL of 7-AAD (559925, BD Pharmingen) at room temperature for 10 min. Subsequently, B cells were examined by flow cytometry to determine the proportion of 7-AAD^+^ dead cells.

### In vitro differentiation of germinal center B cells

WT or Cmtm6 KO primary spleen B cells were cultured in RPMI1640 medium containing 1% P/S, 10% FBS, 50 ng/mL of IL-2, 10 ng/mL of IL-4, 10 ng/mL of IL-21, and 10 ng/mL of BAFF to maintain their survival. To induce the differentiation of primary B cells to germinal center B cells, 2 μg/mL or 5 μg/mL of CD40 agonist was added to the medium and incubated for 72 h. CD95^+^ GL7^+^ germinal center B cells were quantified by flow cytometry.

### T/B cells co-culture

OT-II CD4^+^ T cells were isolated from the spleens of OT-II mice by immunomagnetic negative selection (Mouse CD4^+^ T Cell Isolation Kit, 19852, Stemcell). WT or Cmtm6 KO primary spleen B cells were isolated, grown at 37°C, and divided into two groups. One group was treated for two hours with 25 μg/mL OVA_323-329_ (HY-P0286, MCE), and the other group received PBS treatment. Treated B cells and OT-II CD4^+^ T cells were cultured in 24-well plates at a ratio of 0.5:1 and 2:1. When studying the role of CD40 signaling in T/B interactions, 50 μg/mL of CD40L blocking antibody (BE0017, BioXcell) was added. After 24 h of culture, cells were collected and CD69 and CD25 expressed by CD4^+^ T cells were detected by flow cytometry.

After 72 h of culture, cells were collected and Ki67 expressed by CD4^+^ T cells was detected by flow cytometry. The Transcription Factor Buffer Set (562725, BD Pharmingen) is required for the detection of Ki67.

### Detection of phosphorylated proteins by flow cytometry

2×10^6^ of WT or Cmtm6 KO primary spleen B cells were added with 5 μg/mL of CD40 agonist at 37℃ for 15 min. After centrifugation, 100 μL of Cytofix Fixation buffer (554655, BD Pharmingen) was added to the cell pellet to fix the cells at room temperature for 8 min. After centrifugation, 200 μL of Perm Buffer III (558050, BD Pharmingen) was added to the cell pellet for permeabilization at 4℃ for 30 min. After centrifugation, primary antibodies against phosphorylated proteins were added to the cell pellet at room temperature for 45 min.

Subsequently, the respective fluorescent secondary antibodies were added at room temperature for 45 min. Finally, the levels of phosphorylated proteins were detected by flow cytometry.

### B cell depletion study

Mice were randomly divided and injected with tumor cells (CT26, B16F10 or MC38) subcutaneously, which was defined as day 0. Generally, B cells were deleted by intraperitoneal (i.p.) injection of anti-CD20 (BE0356, BioXcell) on day −2 (150 μg/each) and day 2 (150 μg/each). For the adoptive transfusion of B cells experiments, B cells were deleted by i.p. injection of anti-CD20 on day −12 (150 μg/each) and day −10 (150 μg/each). The deletion effect of CD45^+^ B220^+^ B cells in tumors and dLNs was identified by flow cytometry.

### In vivo tumor models

For all the subcutaneous models, mice were randomly grouped and injected subcutaneously with 1×10^6^ tumor cells (CT26, MC38, B16F10, MC38-OVA or B16-OVA) in the right forelimb.

Tumor volume was measured and recorded every two or three days. Tumor volume was calculated as tumor volume (mm^3^) = tumor length × width × width/2, and tumor growth curves were plotted. At least four time points after tumor inoculation, the mice were sacrificed and the tumors harvested for weighing, photographing, or other purposes. The euthanasia endpoints were set as tumors larger than 2000 mm^3^ or larger than 20 mm in length or mice with significant weight loss.

### In vivo treatment studies

Studies of anti-CD40 (BE0016-2, BioXcell), anti-PD-1 (BE0273, BioXcell) or anti-CD40L (BE0017, BioXcell) therapy in mice bearing CT26, MC38 or B16F10 tumors are consistent with the descriptions above. The administration methods of each drug are shown in the respective flow charts in the figures.

### Immunophenotype analysis

For immunotyping of intratumoral cells, tumor tissues were digested by 1 mg/mL of collagenase IV (40510ES60, Yeasen) and 1 mg/mL of hyaluronidase (20426ES60, Yeasen) and were filtered through 75-micron nylon mesh (7061011, Dakewe) into single cell suspensions and then were subjected to erythrocyte removal by red blood cell lysis buffer (40401ES60, Yeasen). The number of cells in all samples was examined and adjusted to the same cell density.

Then the cells were blocked with 4% FBS and anti-CD16/CD32 (553141, BD Biosciences), incubated with surface marker antibodies for 20 minutes and live/dead fluorescent dye (7-AAD or FVS575V) at 4℃ and then permeabilized with BD Cytofix/Cytoperm buffer (554714) before intracellular labeling antibodies were added for 30 minutes at 4℃.

For immunotyping of splenocytes and lymph node cells, single-cell suspensions were obtained by grinding tissues through the rubber end of a 2 mL syringe and then were filtered through 75-micron nylon mesh. Single cells were subject to erythrocyte removal by red blood cell lysis buffer. The number of cells in all samples was examined and adjusted to the same cell density. The other staining steps are the same as above.

Flow cytometry analysis was performed using ACEA NovoCyte (Agilent) and data processing was done through NovoExpress software (version 1.6.0). Instruments are tested by QC microspheres prior to use. Antibody staining was performed following the manufacturer’s recommendations. Please refer to the key resources table for information about the antibodies used.

### Adoptive transfusion of B cells

Mice were randomly divided and injected with tumor cells (B16F10 or MC38) subcutaneously, which was defined as day 0. B cells were deleted by i.p. injection of anti-CD20 on day −12 (150 μg/each) and day −10 (150 μg/each). When the tumor volume reached approximately 100 mm^3^ (MC38: day 8; B16F10: day 5), 6×10^6^ B cells were injected peritumorally. B cells were obtained from spleens of WT mice or *Cmtm6* KO mice by magnetic cell sorting. Tumor growth kinetics, volumes and weights were measured.

### Single-cell sequencing data analysis

Single cell RNA-seq data used in this study were all from publicly available data. Data for CMTM6 expression in immune cells were obtained from the GEO database (GSE127465). The SpringViewer interactive tool was used to analyze the data.

### RNA-seq

About 30 mg of fresh spleen tissues or 2×10^6^ splenic B cells were obtained from sacrificed mice. RNA isolation, transcriptome libraries construction, sequencing and basic data analysis were conducted by BGI. Based on the RNA-seq raw data, differential expression was evaluated with DESeq. A fold-change of 2:1 or greater and a false discovery rate (FDR)-corrected p-value of 0.05 or less were set as the threshold for differential genes.

### CMTM6-CD40 interaction analysis

For co-immunoprecipitation experiments, we mainly used Immunoprecipitation Kit with Protein G Magnetic Beads (P2177S, Beyotime). Experiments were performed following the manufacturer’s recommendations. Cells were lysed in Lysis Buffer with Protease Inhibitors for 30 min at 4℃. After sufficient lysis, centrifuge the cells at 10,000×g for 5minutes at 4℃ and take the supernatant. The Protein G magnetic beads were mixed with primary antibody and incubated for 1 h at room temperature. The magnetic bead-antibody was then added to the sample lysate, mixed, and incubated at room temperature for 2 h. After three washes in Lysis

Buffer with Protease Inhibitors, samples were eluted in SDS-PAGE Sample Loading Buffer (1×). Finally, the supernatant was used in the Western blot.

### Electrotransfection of hPBMC B cells

Human PBMCs were purchased from Milestone (PB009C-1). The cells were transfected with 200 nM control siRNA or CMTM6 siRNA (5’-CCTTTCTTCTGAGTCTCCTTATACT-3’) using the Neon NxT Electroporation System (Thermo Fisher) according to the manufacturer’s protocol. The siRNA was added to 100 μL of cell suspension at 1×10^^7^ cells/mL in T buffer and electroporated using 1 pulses of 20 ms at 2250 V. After 24 hours of culture, hPBMC B cells were assayed for CMTM6 and CD40 levels by flow cytometry.

### Membranal CD40 degradation assay

Cas9 control A20 cells and CMTM6 KO A20 cells were resuspended in 200 μL of PBS and stained with 8 μL of anti-CD40 PE antibody for 1 h. To terminate the staining, 1 mL of complete medium was added. After washing, 2×10^5^ cells were resuspended with complete medium and cultured in 24-well plates. 50 μM MG132 (HY-13259, MCE) and 50 μM CLQ (HY-17589, MCE) were added separately. Subsequently, cells were collected at 0 h, 2 h, 4 h, and 6 h, and the PE signal was detected by flow cytometry.

### CHX chase assay

2×10^5^ Cas9 control A20 cells and CMTM6 KO A20 cells were resuspended with complete medium and cultured in 24-well plates. 50 μM MG132 (HY-13259, MCE), 50 μM CLQ (HY-17589, MCE), 50 μM ConA (HY-N1724, MCE), and 50 μg/mL of CHX (HY-12320, MCE) were added separately. Subsequently, cells were collected at 6 h and stained with anti-CD40 PE antibody for 30 min. The PE signal was detected by flow cytometry.

### Structure prediction of protein complex

Protein amino acid sequences were obtained from the Uniprot database. AlphaFold2 was utilized to predict the structures of human and mouse CMTM6/CD40 complexes, human and mouse CMTM6/PD-L1 complexes, and human CMTM6/CD58 complexes. We used the Alphafold2-multimer algorithm via Google Colab ^41^ (https://colab.research.google.com/github/sokrypton/ColabFold/blob/main/AlphaFold2.ipynb) to make predictions. Structural presentation of protein complexes and analysis of interaction sites was performed by Discovery studio. The default parameters of each software were used in the calculations.

## QUANTIFICATION AND STATISTICAL ANALYSIS

### Statistical methods and software

The animal experiments were randomized but the operators were not blinded to allocation during experiments and outcome analysis. Statistical analysis was performed using GraphPad Prism 8 Software. No statistical methods were used to predetermine the sample size. A Student’s t test was used for comparison between the two groups. Multiple comparisons were performed using one-way ANOVA followed by Tukey’s multiple comparisons test or two-way ANOVA followed by Tukey’s multiple comparisons test. Detailed statistical methods and sample sizes in the experiments are described in each figure legend. All statistical tests were two-sided and P-values < 0.05 were considered to be significant. ns not significant; *p < 0.05; **p < 0.01; ***p < 0.001.

## REFERENCES

1. Binnewies, M., Roberts, E.W., Kersten, K., Chan, V., Fearon, D.F., Merad, M., Coussens, L.M., Gabrilovich, D.I., Ostrand-Rosenberg, S., Hedrick, C.C., et al. (2018). Understanding the tumor immune microenvironment (TIME) for effective therapy. Nat. Med. 24, 541–550. 10.1038/s41591-018-0014-x.

2. Downs-Canner, S.M., Meier, J., Vincent, B.G., and Serody, J.S. (2022). B Cell Function in the Tumor Microenvironment. Annu Rev Immunol 40, 169–193. 10.1146/annurev-immunol-101220-015603.

3. DiLillo, D.J., Yanaba, K., and Tedder, T.F. (2010). B cells are required for optimal CD4+ and CD8+ T cell tumor immunity: therapeutic B cell depletion enhances B16 melanoma growth in mice. J Immunol 184, 4006–4016. 10.4049/jimmunol.0903009.

4. Cui, C., Wang, J., Fagerberg, E., Chen, P.M., Connolly, K.A., Damo, M., Cheung, J.F., Mao, T., Askari, A.S., Chen, S., et al. (2021). Neoantigen-driven B cell and CD4 T follicular helper cell collaboration promotes anti-tumor CD8 T cell responses. Cell 184, 6101–6118 e6113. 10.1016/j.cell.2021.11.007.

5. Cabrita, R., Lauss, M., Sanna, A., Donia, M., Skaarup Larsen, M., Mitra, S., Johansson, I., Phung, B., Harbst, K., Vallon-Christersson, J., et al. (2020). Tertiary lymphoid structures improve immunotherapy and survival in melanoma. Nature 577, 561–565. 10.1038/s41586-019-1914-8.

6. Helmink, B.A., Reddy, S.M., Gao, J., Zhang, S., Basar, R., Thakur, R., Yizhak, K., Sade-Feldman, M., Blando, J., Han, G., et al. (2020). B cells and tertiary lymphoid structures promote immunotherapy response. Nature 577, 549–555. 10.1038/s41586-019-1922-8.

7. Petitprez, F., de Reynies, A., Keung, E.Z., Chen, T.W., Sun, C.M., Calderaro, J., Jeng, Y.M., Hsiao, L.P., Lacroix, L., Bougouin, A., et al. (2020). B cells are associated with survival and immunotherapy response in sarcoma. Nature 577, 556–560. 10.1038/s41586-019-1906-8.

8. Hao, D., Han, G., Sinjab, A., Gomez-Bolanos, L.I., Lazcano, R., Serrano, A., Hernandez, S.D., Dai, E., Cao, X., Hu, J., et al. (2022). The Single-Cell Immunogenomic Landscape of B and Plasma Cells in Early-Stage Lung Adenocarcinoma. Cancer Discov. 12, 2626–2645. 10.1158/2159-8290.CD-21-1658.

9. Bod, L., Kye, Y.C., Shi, J., Torlai Triglia, E., Schnell, A., Fessler, J., Ostrowski, S.M., Von-Franque, M.Y., Kuchroo, J.R., Barilla, R.M., et al. (2023). B-cell-specific checkpoint molecules that regulate anti-tumour immunity. Nature 619, 348–356. 10.1038/s41586-023-06231-0.

10. Zhang, Z., Xu, X., Zhang, D., Zhao, S., Wang, C., Zhang, G., Chen, W., Liu, J., Gong, H., Rixiati, Y., et al. (2024). Targeting Erbin-mitochondria axis in platelets/megakaryocytes promotes B cell-mediated antitumor immunity. Cell Metab. 10.1016/j.cmet.2023.12.020.

11. Han, W., Ding, P., Xu, M., Wang, L., Rui, M., Shi, S., Liu, Y., Zheng, Y., Chen, Y., Yang, T., and Ma, D. (2003). Identification of eight genes encoding chemokine-like factor superfamily members 1–8 (CKLFSF1–8) by in silico cloning and experimental validation. Genomics 81, 609–617. 10.1016/s0888-7543(03)00095-8.

12. Liang, J., Li, S., Li, W., Rao, W., Xu, S., Meng, H., Zhu, F., Zhai, D., Cui, M., Xu, D., et al. (2022). CMTM6, a potential immunotherapy target. J. Cancer Res. Clin. Oncol. 148, 47-56. 10.1007/s00432-021-03835-9.

13. Yaseen, M.M., Abuharfeil, N.M., and Darmani, H. (2022). CMTM6 as a master regulator of PD-L1. Cancer Immunol. Immunother. 10.1007/s00262-022-03171-y.

14. Liu, Q., Wang, J., Guo, Z., Zhang, H., Zhou, Y., Wang, P., Li, T., Lu, W., Liu, F., and Han, W. (2023). CMTM6 promotes hepatocellular carcinoma progression through stabilizing beta-catenin. Cancer Lett, 216585. 10.1016/j.canlet.2023.216585.

15. Mohapatra, P., Shriwas, O., Mohanty, S., Ghosh, A., Smita, S., Kaushik, S.R., Arya, R., Rath, R., Das Majumdar, S.K., Muduly, D.K., et al. (2021). CMTM6 drives cisplatin resistance by regulating Wnt signaling through the ENO-1/AKT/GSK3beta axis. JCI Insight 6, e143643. 10.1172/jci.insight.143643.

16. Burr, M.L., Sparbier, C.E., Chan, Y.C., Williamson, J.C., Woods, K., Beavis, P.A., Lam, E.Y.N., Henderson, M.A., Bell, C.C., Stolzenburg, S., et al. (2017). CMTM6 maintains the expression of PD-L1 and regulates anti-tumour immunity. Nature 549, 101–105. 10.1038/nature23643.

17. Mezzadra, R., Sun, C., Jae, L.T., Gomez-Eerland, R., de Vries, E., Wu, W., Logtenberg, M.E.W., Slagter, M., Rozeman, E.A., Hofland, I., et al. (2017). Identification of CMTM6 and CMTM4 as PD-L1 protein regulators. Nature 549, 106–110. 10.1038/nature23669.

18. Ho, P., Melms, J.C., Rogava, M., Frangieh, C.J., Pozniak, J., Shah, S.B., Walsh, Z., Kyrysyuk, O., Amin, A.D., Caprio, L., et al. (2023). The CD58-CD2 axis is co-regulated with PD-L1 via CMTM6 and shapes anti-tumor immunity. Cancer Cell 41, 1207–1221 e1212. 10.1016/j.ccell.2023.05.014.

19. Miao, B., Hu, Z., Mezzadra, R., Hoeijmakers, L., Fauster, A., Du, S., Yang, Z., Sator-Schmitt, M., Engel, H., Li, X., et al. (2023). CMTM6 shapes antitumor T cell response through modulating protein expression of CD58 and PD-L1. Cancer Cell 41, 1817–1828 e1819. 10.1016/j.ccell.2023.08.008.

20. Jia, X.M., Long, Y.R., Yu, X.L., Chen, R.Q., Gong, L.K., and Geng, Y. (2022). Construction of stable membranal CMTM6-PD-L1 full-length complex to evaluate the PD-1/PD-L1-CMTM6 interaction and develop anti-tumor anti-CMTM6 nanobody. Acta Pharmacol. Sin. 44, 1095–1104. 10.1038/s41401-022-01020-3.

21. Long, Y., Chen, R., Yu, X., Tong, Y., Peng, X., Li, F., Hu, C., Sun, J., and Gong, L. (2023). Suppression of tumor or host intrinsic CMTM6 drives antitumor cytotoxicity in a PD-L1-independent manner. Cancer Immunol Res 11, 241–260. 10.1158/2326-6066.CIR-22-0439.

22. Zilionis, R., Engblom, C., Pfirschke, C., Savova, V., Zemmour, D., Saatcioglu, H.D., Krishnan, I., Maroni, G., Meyerovitz, C.V., Kerwin, C.M., et al. (2019). Single-Cell Transcriptomics of Human and Mouse Lung Cancers Reveals Conserved Myeloid Populations across Individuals and Species. Immunity 50, 1317–1334.e1310. 10.1016/j.immuni.2019.03.009.

23. Qiao, G., Lei, M., Li, Z., Sun, Y., Minto, A., Fu, Y.X., Ying, H., Quigg, R.J., and Zhang, J. (2007). Negative regulation of CD40-mediated B cell responses by E3 ubiquitin ligase Casitas-B-lineage lymphoma protein-B. J Immunol 179, 4473–4479. 10.4049/jimmunol.179.7.4473.

24. Rowland, S.L., Tremblay, M.M., Ellison, J.M., Stunz, L.L., Bishop, G.A., and Hostager, B.S. (2007). A novel mechanism for TNFR-associated factor 6-dependent CD40 signaling. J Immunol 179, 4645–4653. 10.4049/jimmunol.179.7.4645.

25. Hostager, B.S., Catlett, I.M., and Bishop, G.A. (2000). Recruitment of CD40 and tumor necrosis factor receptor-associated factors 2 and 3 to membrane microdomains during CD40 signaling. J Biol Chem 275, 15392–15398. 10.1074/jbc.M909520199.

26. Bandyopadhyay, S., Gurjar, D., Saha, B., and Bodhale, N. (2023). Decoding the contextual duality of CD40 functions. Hum. Immunol. 84, 590–599. 10.1016/j.humimm.2023.08.142.

27. Qin, Z., Richter, G., Schüler, T., Ibe, S., Cao, X., and Blankenstein, T. (1998). B cells inhibit induction of T cell-dependent tumor immunity. Nat. Med. 4, 627–630. 10.1038/nm0598-627.

28. Perricone, M.A., Smith, K.A., Claussen, K.A., Plog, M.S., Hempel, D.M., Roberts, B.L., St George, J.A., and Kaplan, J.M. (2004). Enhanced efficacy of melanoma vaccines in the absence of B lymphocytes. J. Immunother. 27, 273–281. 10.1097/00002371-200407000-00003.

29. Mizoguchi, A., Mizoguchi, E., Takedatsu, H., Blumberg, R.S., and Bhan, A.K. (2002). Chronic intestinal inflammatory condition generates IL-10-producing regulatory B cell subset characterized by CD1d upregulation. Immunity 16, 219–230. 10.1016/s1074-7613(02)00274-1.

30. Pylayeva-Gupta, Y., Das, S., Handler, J.S., Hajdu, C.H., Coffre, M., Koralov, S.B., and Bar-Sagi, D. (2016). IL35-Producing B Cells Promote the Development of Pancreatic Neoplasia. Cancer Discov. 6, 247–255. 10.1158/2159-8290.CD-15-0843.

31. Takahashi, R., Macchini, M., Sunagawa, M., Jiang, Z., Tanaka, T., Valenti, G., Renz, B.W., White, R.A., Hayakawa, Y., Westphalen, C.B., et al. (2021). Interleukin-1β-induced pancreatitis promotes pancreatic ductal adenocarcinoma via B lymphocyte-mediated immune suppression. Gut 70, 330–341. 10.1136/gutjnl-2019-319912.

32. Xiao, X., Lao, X.M., Chen, M.M., Liu, R.X., Wei, Y., Ouyang, F.Z., Chen, D.P., Zhao, X.Y., Zhao, Q., Li, X.F., et al. (2016). PD-1hi Identifies a Novel Regulatory B-cell Population in Human Hepatoma That Promotes Disease Progression. Cancer Discov. 6, 546–559. 10.1158/2159-8290.CD-15-1408.

33. Shalapour, S., Lin, X.J., Bastian, I.N., Brain, J., Burt, A.D., Aksenov, A.A., Vrbanac, A.F., Li, W., Perkins, A., Matsutani, T., et al. (2017). Inflammation-induced IgA+ cells dismantle anti-liver cancer immunity. Nature 551, 340–345. 10.1038/nature24302.

34. Biswas, S., Mandal, G., Payne, K.K., Anadon, C.M., Gatenbee, C.D., Chaurio, R.A., Costich, T.L., Moran, C., Harro, C.M., Rigolizzo, K.E., et al. (2021). IgA transcytosis and antigen recognition govern ovarian cancer immunity. Nature 591, 464–470. 10.1038/s41586-020-03144-0.

35. Tang, T., Cheng, X., Truong, B., Sun, L., Yang, X., and Wang, H. (2021). Molecular basis and therapeutic implications of CD40/CD40L immune checkpoint. Pharmacol. Ther. 219, 107709. 10.1016/j.pharmthera.2020.107709.

36. Xu, J., Foy, T.M., Laman, J.D., Elliott, E.A., Dunn, J.J., Waldschmidt, T.J., Elsemore, J., Noelle, R.J., and Flavell, R.A. (1994). Mice deficient for the CD40 ligand. Immunity 1, 423–431. 10.1016/1074-7613(94)90073-6.

37. Bergqvist, P., Gardby, E., Stensson, A., Bemark, M., and Lycke, N.Y. (2006). Gut IgA class switch recombination in the absence of CD40 does not occur in the lamina propria and is independent of germinal centers. J Immunol 177, 7772–7783. 10.4049/jimmunol.177.11.7772.

38. Chen, Y., Yu, M., Zheng, Y., Fu, G., Xin, G., Zhu, W., Luo, L., Burns, R., Li, Q.Z., Dent, A.L., et al. (2019). CXCR5(+)PD-1(+) follicular helper CD8 T cells control B cell tolerance. Nat Commun 10, 4415. 10.1038/s41467-019-12446-5.

39. Koike, T., Harada, K., Horiuchi, S., and Kitamura, D. (2019). The quantity of CD40 signaling determines the differentiation of B cells into functionally distinct memory cell subsets. Elife 8. 10.7554/eLife.44245.

40. Xing, F., Gao, H., Chen, G., Sun, L., Sun, J., Qiao, X., Xue, J., and Liu, C. (2023). CMTM6 overexpression confers trastuzumab resistance in HER2-positive breast cancer. Mol Cancer 22, 6. 10.1186/s12943-023-01716-y.

41. Mirdita, M., Schutze, K., Moriwaki, Y., Heo, L., Ovchinnikov, S., and Steinegger, M. (2022). ColabFold: making protein folding accessible to all. Nat. Methods 19, 679–682. 10.1038/s41592-022-01488-1.

