## Supplementary material for "CMTM6 maintains B cell-intrinsic CD40 expression to regulate anti-tumor immunity": Main text

**This file contains:**

Figure S1. Effects of B-cell deletion on tumor immunity.

Figure S2. Effects of B-cell deletion on in situ intestinal cancer model.

Figure S3. CMTM6 expression in B cells.

Figure S4. B-cell deletion and adoptive transfer.

Figure S5. Identification of B cell CMTM6 knockout mice.

Figure S6. Effects of B cell CMTM6 deficiency on T/B cell interaction.

Figure S7. Flow-cytometry gating strategy for B cell subsets.

Figure S8. B cell differentiation in CMTM6 knockout mice.

Figure S9. Activation of CMTM6 knockout B cells by several agonists.

Figure S10. Activation of splenic B cells by CD40 agonist.

Figure S11. Multi-omics data mining.

Figure S12. CMTM6 knockdown in B cells of human PBMCs affects CD40 levels.

Figure S13. Structure prediction of CMTM6 protein complexes by AlphaFold2.

### Supplementary figures

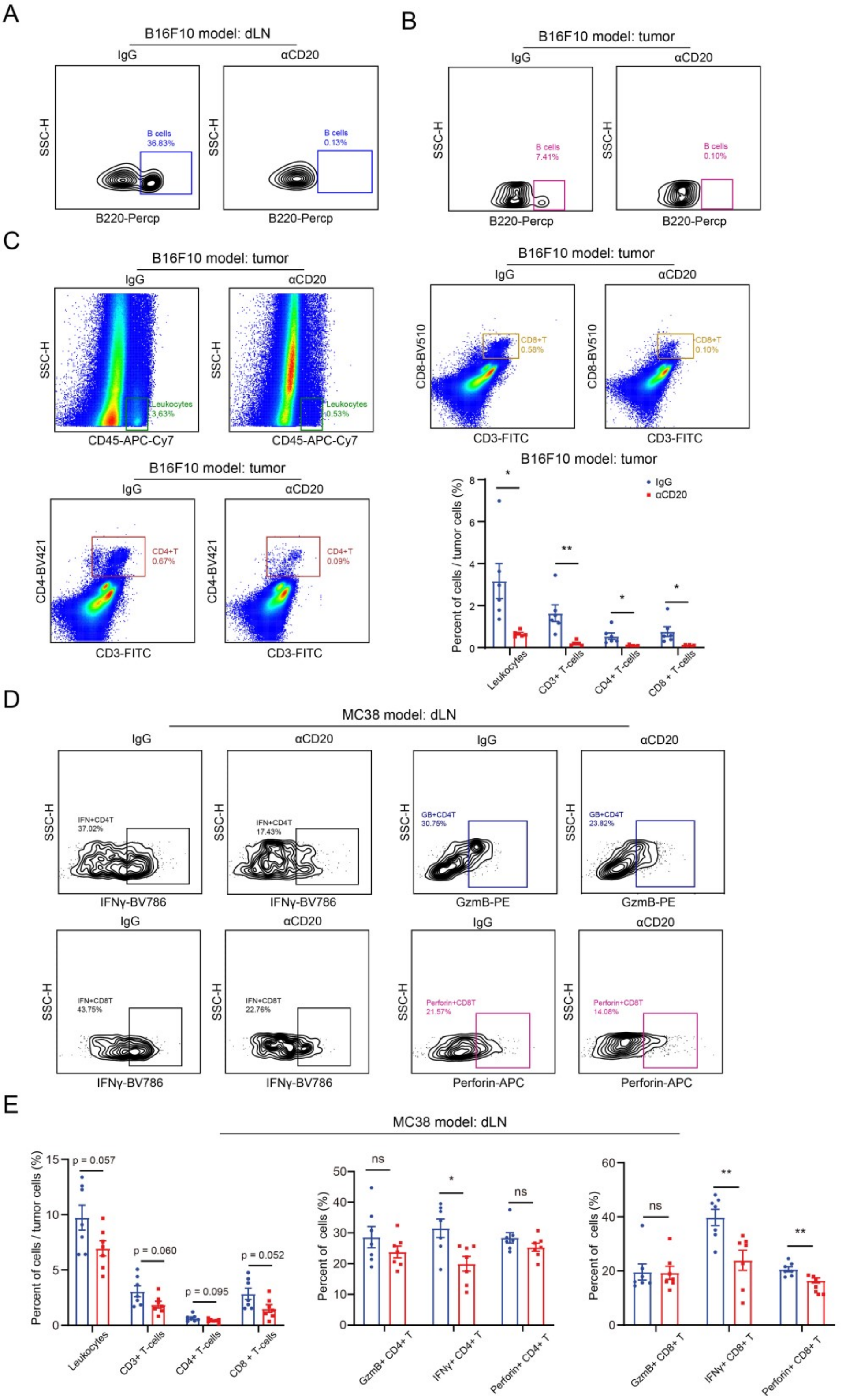


**Figure S1. Effects of B-cell deletion on tumor immunity.**

(A and B) Validation of the effect of B-cell deletion in dLNs (A) and tumors (B) by flow cytometry.

(C) Effect of B-cell deletion on intratumoral infiltration of leukocytes and T cells in the B16F10 tumor model (n = 7).

(D and E) Effect of B-cell deletion on effector molecule levels of dLN T cells in the MC38 tumor model (n = 7).

The data are presented as the mean ± SEM. * $p < 0.05$; ** $p < 0.01$; ns not significant by two-way ANOVA followed by Tukey's multiple comparisons test.

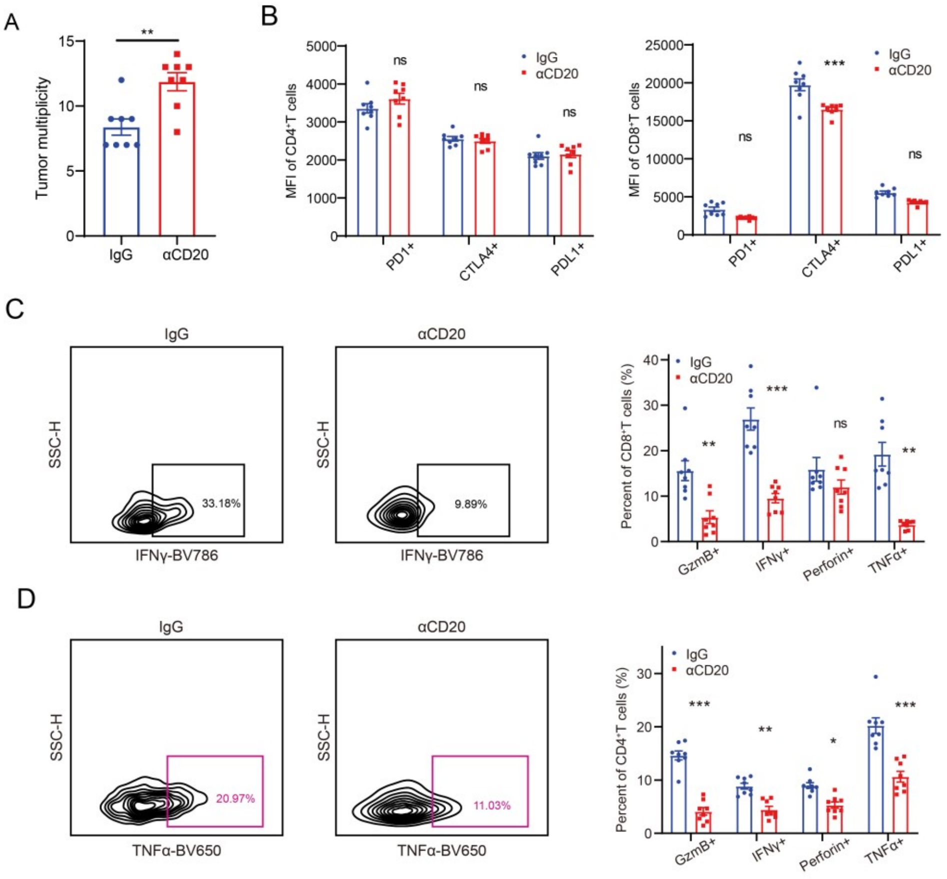


**Figure S2. Effects of B-cell deletion on in situ intestinal cancer model.**

(A) Effect of B-cell deletion on tumor multiplicity in the APC-min model (n = 8).

(B) Effect of B-cell deletion on inhibitory molecule levels of MLN T cells in the APC-min model (n = 8).

(C) Effect of B-cell deletion on effector molecule levels of MLN $CD8^+$ T cells in the APC-min model (n = 8).

(D) Effect of B-cell deletion on effector molecule levels of MLN $CD4^+$ T cells in the APC-min model (n = 8).

The data are presented as the mean ± SEM. * $p < 0.05$; ** $p < 0.01$; *** $p < 0.001$; ns not significant by unpaired t test or two-way ANOVA followed by Tukey’s multiple comparisons test.

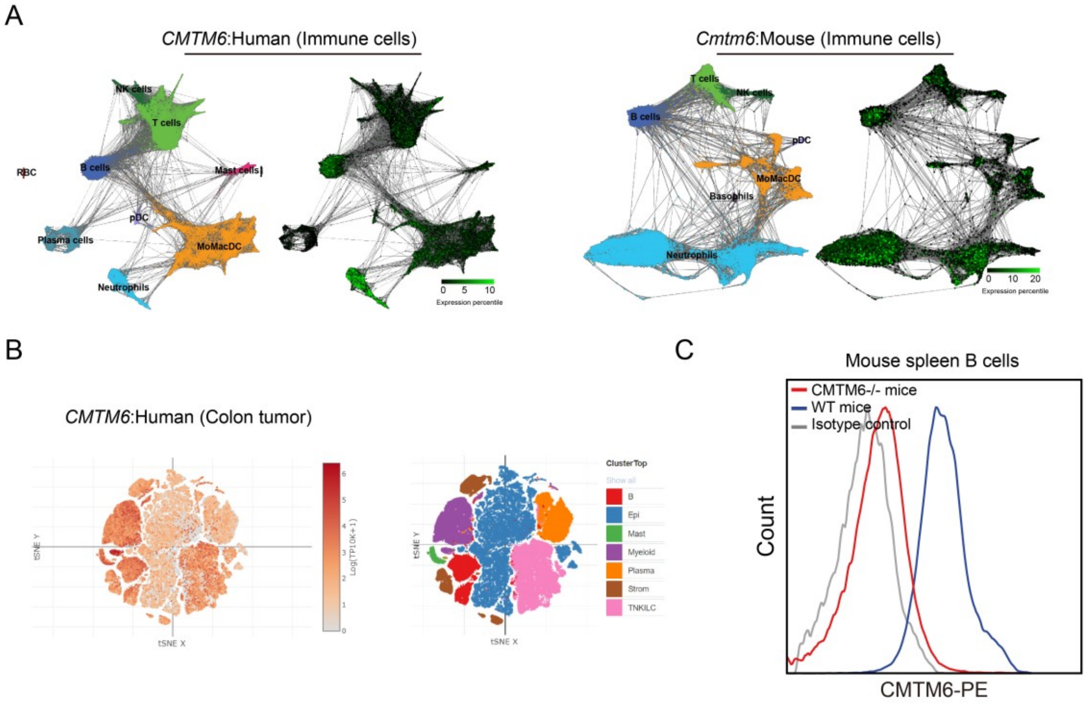


**Figure S3. CMTM6 expression in B cells.**

(A) CMTM6 expression analysis by single-cell sequencing of human and mouse CD45$^{+}$ immune cells.

(B) CMTM6 expression analysis by single-cell sequencing of open-source clinical human colon cancer samples.

(C) CMTM6 expression in splenic B cells of WT and *Cmtm6* KO mice was analyzed by flow cytometry.

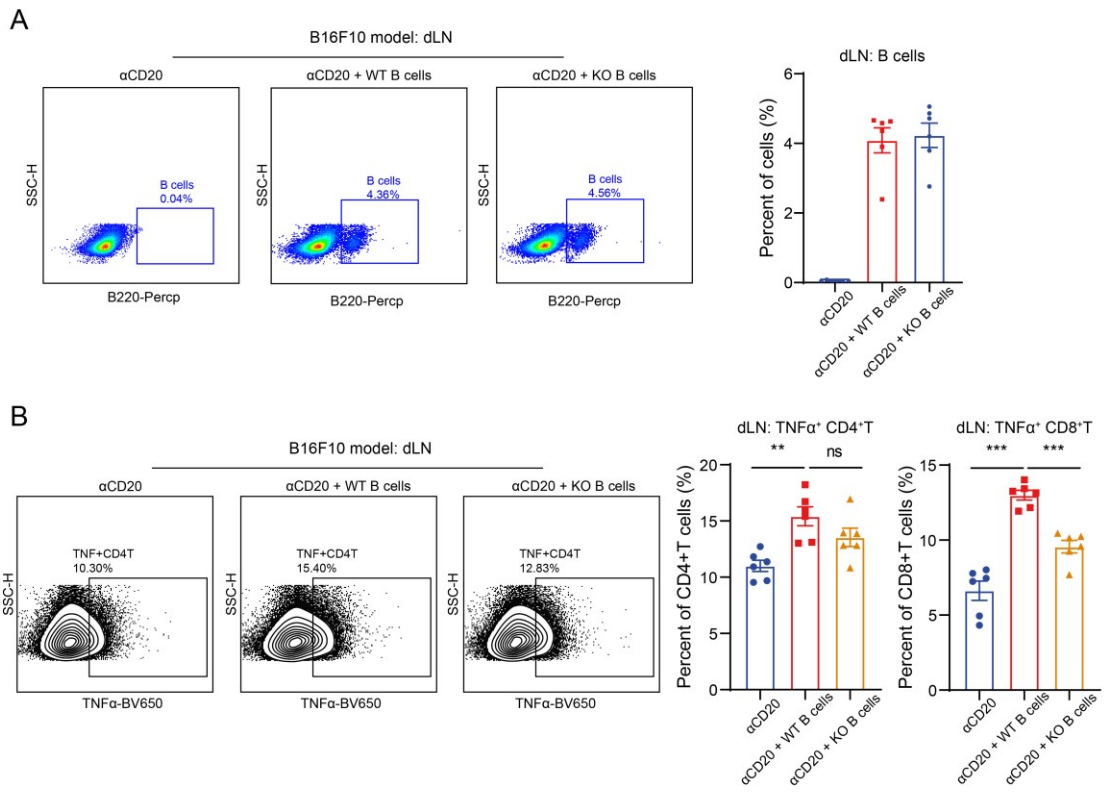


**Figure S4. B-cell deletion and adoptive transfer.**

(A) Validation of the effect of B-cell deletion and B-cell transfusion back in the B16F10 tumor model (n = 6).

(B) Effector molecule analysis of $CD4^+$ T cells and $CD8^+$ T cells in dLNs of the B16F10 tumor model after WT or *Cmtm6* KO B cells were transfused back (n = 6).

The data are presented as the mean ± SEM. ** $p < 0.01$; ** $p < 0.001$; ns not significant by one-way ANOVA followed by Tukey's multiple comparisons test.

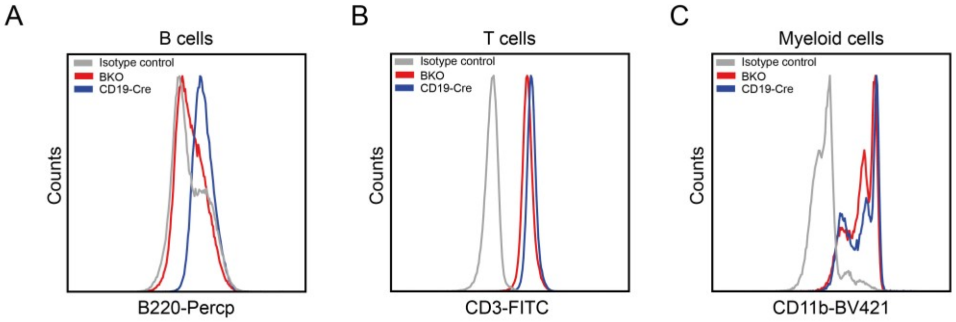


**Figure S5. Identification of B cell CMTM6 knockout mice.**

Representative histogram plots for CMTM6 expression in B cells, T cells, and myeloid cells of CD19-iCre or BKO mice.

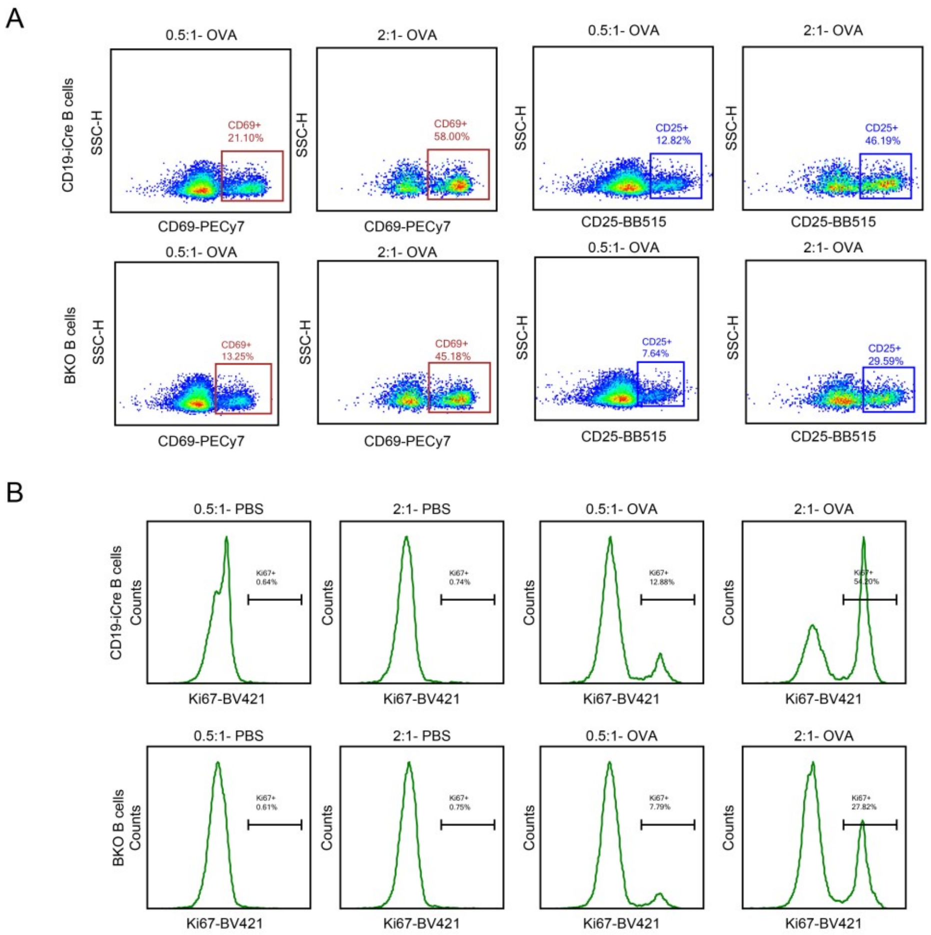


**Figure S6. Effects of B cell CMTM6 deficiency on T/B cell interaction.**

(A) Representative density plots for OT-II CD4$^{+}$ T cell activation when co-cultured with CD19-iCre or BKO B cells for 24 h.

(B) Representative histogram plots for OT-II CD4$^{+}$ T cell proliferation when co-cultured with CD19-iCre or BKO B cells for 72 h.

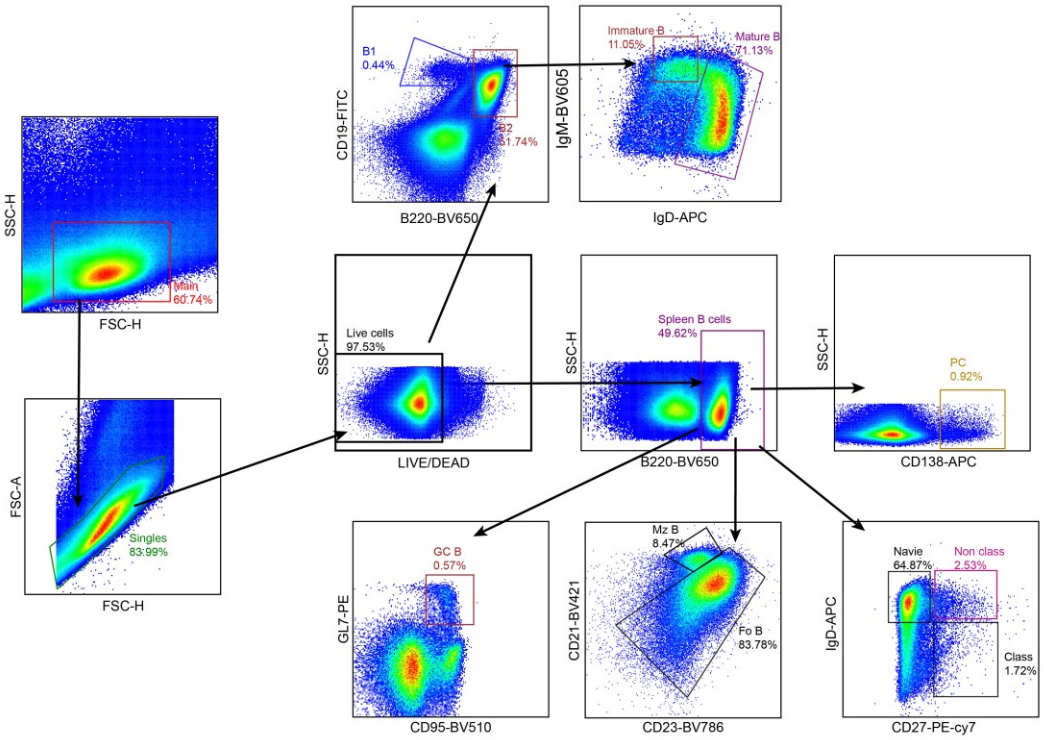


**Figure S7. Flow-cytometry gating strategy for B cell subsets.** Representative flow-cytometry gating strategy for quantifying the B1 cells, B2 cells, marginal zone B cells (Mz B), follicular B cells (Fo B), immature B cells, mature B cells, plasma cells (PC), non-class-switching memory B cells, class-switching memory B cells, and germinal center B cells (GC B) in the mouse spleen.

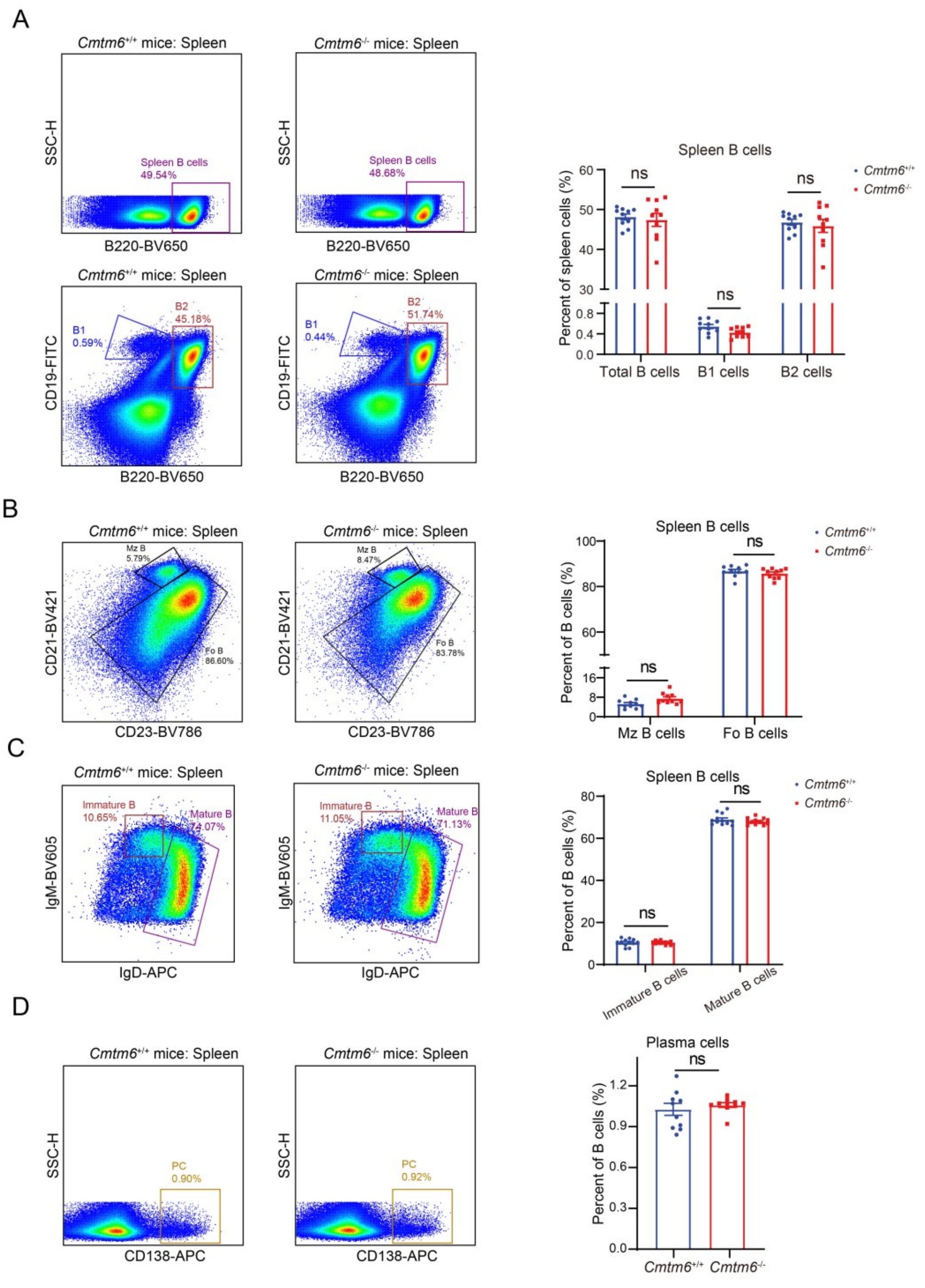


**Figure S8. B cell differentiation in CMTM6 knockout mice.** Representative flow-cytometry gating strategy and statistical analysis of B1 cells, B2 cells, marginal zone B cells (Mz B), follicular B cells (Fo B), immature B cells, mature B cells, and plasma cells (PC). The data are presented as the mean ± SEM. ns not significant by unpaired t test or two-way ANOVA followed by Tukey's multiple comparisons test.

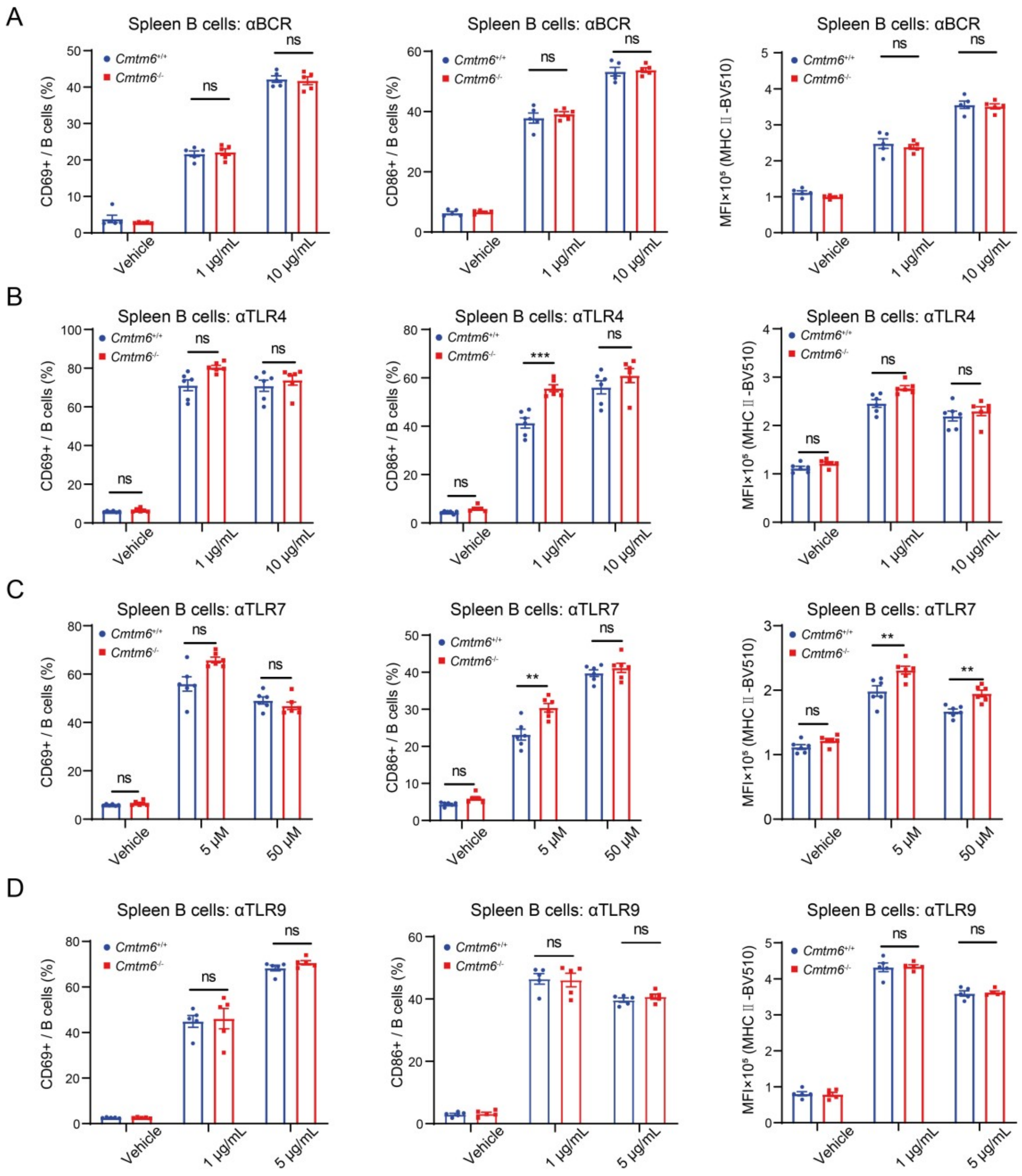


**Figure S9. Activation of CMTM6 knockout B cells by several agonists.** Statistical analysis of the proportion of CD69$^+$, the proportion of CD86$^+$, and the MFI of MHC II in splenic B cells from WT or *Cmtm6* KO mice after the action of BCR agonist (n = 5), TLR4 agonist (n = 6), TLR7 agonist (n = 6), or TLR9 agonist (n = 5). The data are presented as the mean ± SEM. ** p < 0.01; *** p < 0.001; ns not significant by two-way ANOVA followed by Tukey’s multiple comparisons test.

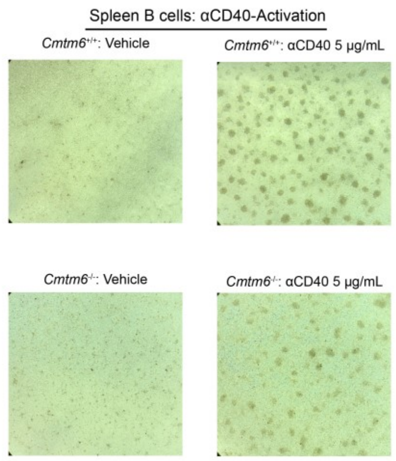


**Figure S10. Activation of splenic B cells by CD40 agonist.** Photographs of sorted WT or *Cmtm6* KO B cells after 24 hours of treatment with PBS or CD40 agonist.

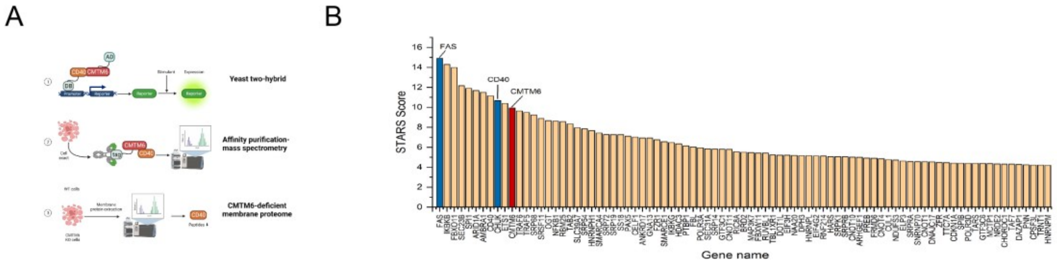


**Figure S11. Multi-omics data mining.**

(A) Open-source yeast two-hybrid and proteomics data mining revealed that CMTM6 interacted with CD40.

(B) Open-source CRISPR screening data mining revealed that CMTM6 KO affected CD40 levels in Daudi cells.

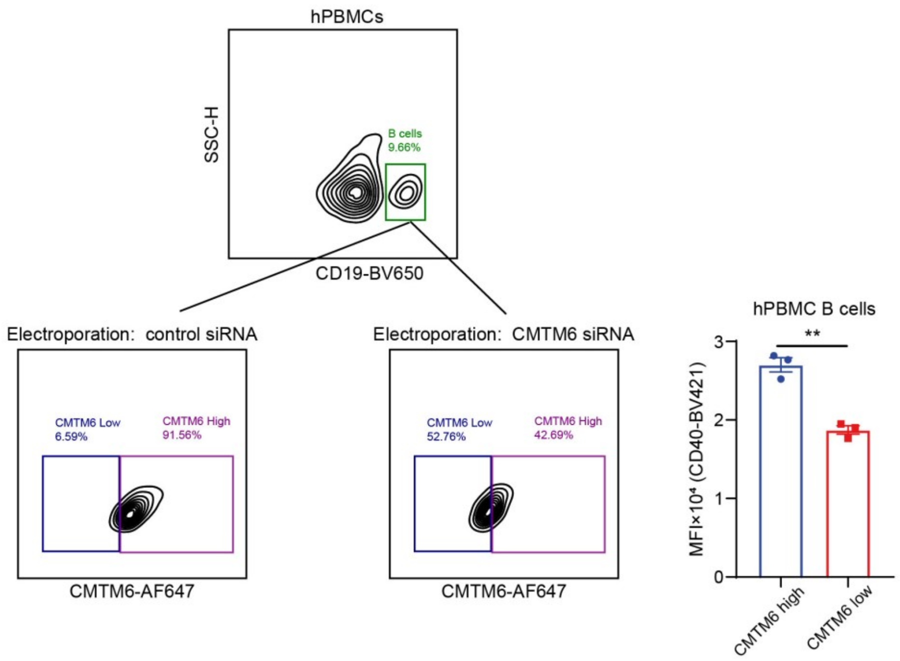


**Figure S12. CMTM6 knockdown in B cells of human PBMCs affects CD40 levels.** Based on siRNA electrotransfection, the levels of CMTM6 and CD40 in human PBMC B cells were analyzed by flow cytometry. The data are presented as the mean ± SEM. ** $p < 0.01$ by unpaired t test.

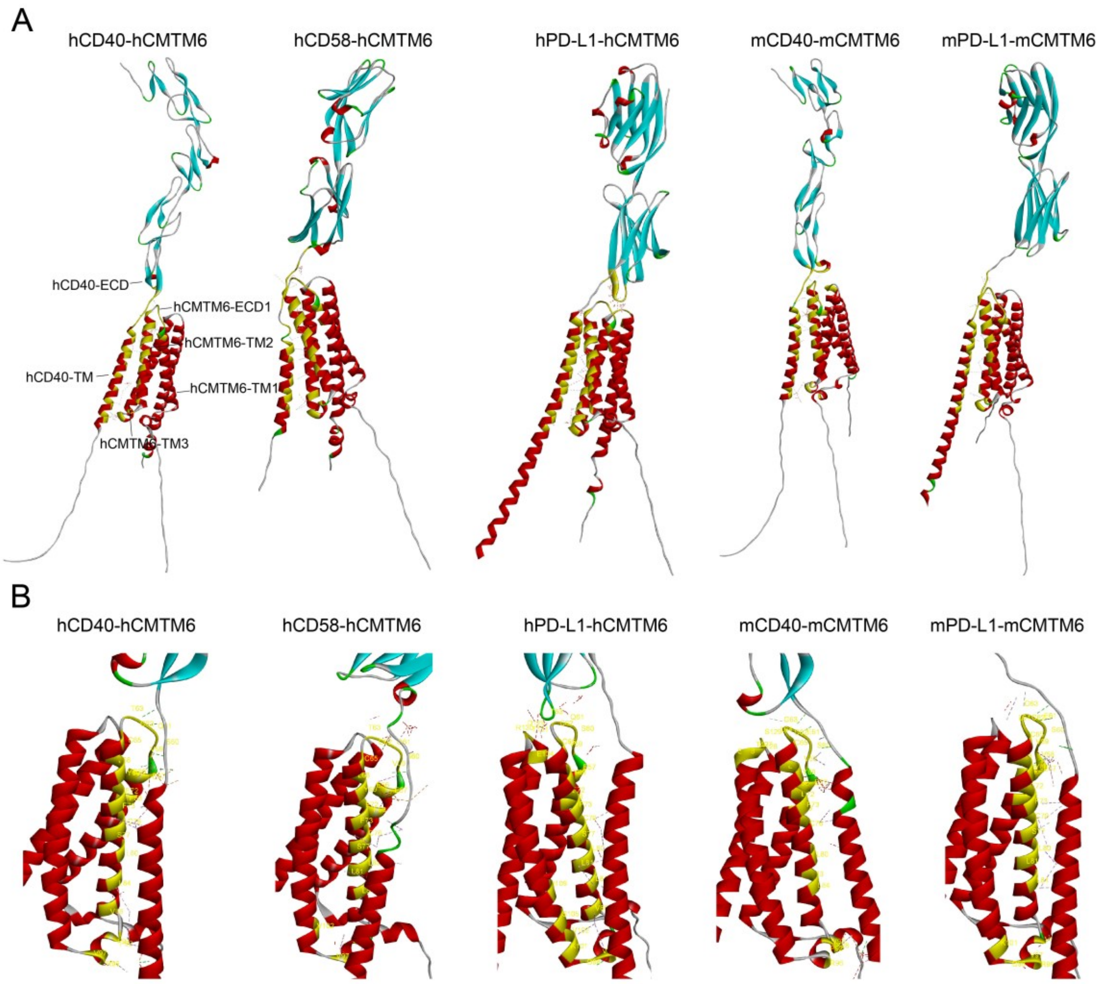


**Figure S13. Structure prediction of CMTM6 protein complexes by AlphaFold2.**

(A) Structure prediction of full-length protein complexes of CMTM6 and PD-L1, CD58, CD40 by AlphaFold2.

(B) Key interaction sites in the predicted structures of CMTM6 with PD-L1, CD58 or CD40.
